# Acoustic features of and behavioral responses to emotionally intense mouse vocalizations

**DOI:** 10.1101/2025.01.12.632636

**Authors:** Zahra Ghasemahmad, Karthic Drishna Perumal, Bhavya Sharma, Rishita Panditi, Jeffrey James Wenstrup

**Author notes:** Indicates equal contributions (Karthic Drishna Perumal, Bhavya Sharma, Rishita Panditi). **Corresponding author:** Jeffrey Wenstrup, Department of Biomedical Sciences, Northeast Ohio Medical University, 4209 State Route 44, Rootstown, Ohio, 44272, USA. Massachusetts Eye and Ear Infirmary, 243 Charles Street, Boston, MA 02114. 5300 North Meadows Drive. Grove City, OH,43123. 1200 Cinnamon Hill Lane, Apt. 201, Columbia, MO, 65201. 100 N. Academy Avenue, Danville, PA 17822.

## Abstract

Social vocalizations contain cues that reflect the motivational state of vocalizing animals. Once perceived, these cues affect the internal state and behavioral responses of listening animals. Using CBA/CAJ mice, this study examined acoustic cues that signal intensity in male-female interactions, then compared behavioral responses to intense mating vocal sequences with those from another intense behavioral context, restraint. Experiment I examined behaviors and vocalizations associated with male-female social interactions. Based on several behaviors, we distinguished more general, courtship-type interactions from mating interactions involving mounting or attempted mounting behaviors. We then compared vocalizations between courtship and mating. The increase in behavioral intensity from courtship to mating was associated with altered syllable composition, more harmonic structure, lower minimum frequency, longer duration, reduced inter-syllable interval, and increased sound intensity. Based on these features, we constructed highly salient playback stimuli associated with mating. In Experiment II, we compared behavioral responses to playback of these mating sequences with responses to playback of aversive vocal sequences produced by restrained mice. Subjects were males and estrus females. We observed a range of behavioral responses. Some (e.g., Attending and Stretch-Attend) showed similar responses across playback type and sex, while others were context dependent (e.g., Flinching, Locomotion). Still other behaviors showed either an effect of sex (e.g., Self-Grooming, Still-and-Alert) or an interaction between playback type and sex (Escape). These results demonstrate both state-dependent features of mouse vocalizations and their effectiveness in evoking a range of behavioral responses, independent of contextual cues provided by other sensory stimuli or behavioral interactions.

## 1. INTRODUCTION

Humans and other species use vocal communication for expressing thoughts, internal state, and emotions during social interactions in order to affect the thoughts, internal states, or behaviors of listeners. Acoustic cues that shape the listener’s responses are generally represented in the segmental features of vocal communication (syllables, words) and suprasegmental features that contribute to prosody, including spectral, temporal, and amplitude cues. Both the choice of syllables and the prosodic features of sounds can be affected by internal state and emotions of the sender [1–4]. Once perceived, these cues may in turn affect the internal state and behavioral responses of the listening animal. Behavioral reactions to salient and valent vocalizations can further be influenced by the listener’s internal state and by previous experience with the vocalizations [5–8]. This study examines acoustic cues that signal intensity in male-female interactions, then compares behavioral responses to intense mating vocal sequences with those from another intense behavioral context, restraint.

In mice, like other species, cues such as syllable types and spectrotemporal characteristics convey information linked to the internal affective state of the sender. Ultrasonic vocalizations (USVs), produced during many social interactions, are modulated in spectrotemporal characteristics in context-dependent ways. In mating and in restraint, intense interactions are associated with lower ultrasonic frequencies [9–12]. Duration is heavily affected by the nature of social interaction: USVs emitted in a mating context are significantly longer than those emitted in restraint or isolation [13,14], and USVs emitted during male fighting are longer than those emitted during fleeing [15]. Vocalizations in mice also include non-USV categories such as low frequency harmonic (LFH) calls (the mouse squeak), mid-frequency vocalizations (MFVs), and Noisy calls that are used for social communication in several behavioral contexts [11,13,15–19]. These non-USVs contain frequencies corresponding to the most sensitive part of the mouse audiogram [20–22] and are likely to travel further distances due to their longer wavelengths. Like USVs, these also show changes in their acoustic features in a context-dependent manner [13,23].

The reception of vocal communication signals has been linked to many behavioral responses and internal state changes. For instance, rodent pups emit vocalizations that trigger maternal/paternal nest building, licking, and crouching behaviors that help nursing [24–26]. These calls, when produced under isolation, cause retrieval behavior in mothers [9,25,27]. During mating behavior, females prefer vocalizing male mice over devocalized males [28,29] and playback of courtship USVs triggers a female’s approach [30]. Further, the patterns of USVs during social interactions can affect the behavior of socially engaged mice but not the behavior of other animals that observe the interaction [15]. Such impact of vocalizations on the listener, both physiologically and behaviorally, is shared by non-USVs [8,31–33].

While our work is informed by the extensive studies on sender vocalizations and listener responses to various vocal categories and acoustic cues, this study seeks specifically to understand how vocalizations with opposing valence, carrying prosodic and semantic information in natural sequences, can influence the behavior of the listener. Experiment I in this study distinguishes the features of vocal sequences that reflect lower and higher intensity of interactions between males and females, in order to create highly salient playback stimuli associated with mating. These are used in Experiment II, which evaluates the behavioral responses of male and female mice to highly emotional vocal sequences associated with mating and with an aversive behavioral context (restraint). We hypothesize that oppositely valent vocal sequences, carrying cues reflecting the internal state of vocalizing animals, can change the behavioral responses in a sex- and context-specific manner.

## 2. MATERIALS AND METHODS

These experiments share several features with our recently published work [8] but differ in other ways. Both studies utilized the same mating and restraint vocal exemplars, presented in the same order, during playback experiments. The present study utilized a larger set of male-female pairs to examine vocal features associated with courtship and mating behaviors. While both studies used the same set of behaviors to assess responses to playback, each study utilized a completely different pool of subjects in playback experiments. Unlike subjects in the previously published work, subjects in this study did not have implanted microdialysis probes during playback.

### 2.1 Animals

Experimental procedures on animal subjects were approved by the Institutional Animal Care and Use Committee at Northeast Ohio Medical University (Protocol 18-09-207). A total of 94 adult CBA/CaJ mice (48 males, 46 females) were used for this study (Table 1). The mice were obtained from The Jackson Laboratory (Bar Harbor, ME) and were aged p90-p180 at the time of experiments. The mice were housed in same-sex groups of 2-4 individuals until the week of the experiments, during which they were singly housed. Cages housing males and females were placed on separate shelves to minimize male-female interactions prior to the experiments. Food and water were provided *ad libitum* except during the experimental sessions that included behavioral recording, experiencing, and playback. Animals were maintained on a reversed dark/light cycle and experimental sessions took place during the dark cycle.

**Table 1:** Animal groups and numbers.

| **Experimental Session** | **Behavior Type** | **Males** | **Females** | **Total** |
| --- | --- | --- | --- | --- |
| **Vocal Recording** | **Mating** | 8 | 8 | 16 |
|  | **Restraint** | 4 | 2 | 6 |
| **Vocal Playback** | **Mating** | 8 | 11 | 19 |
|  | **Restraint** | 8 | 9 | 17 |
| **Experience Only** | **Mating** | 20 | 16 | 36 |
| **Totals** |  | 48 | 46 | 94 |

The estrous stage of female mice was evaluated based on vaginal smear samples obtained by sterile vaginal lavage. Samples were collected using glass pipettes filled with double distilled water, placed on a slide, stained using crystal violet, and coverslipped for microscopic examination. Estrous stage was determined by the predominant cell type: squamous epithelial cells (estrus), nucleated cornified cells (proestrus), or leukocytes (diestrus) [34]. To confirm that the stage of estrous did not change during the experiment day, samples obtained prior to and after data collection on the experimental day were compared.

### 2.2 Structure of experimental sessions and animal groups

Three types of experimental sessions occurred: vocalization recording, experiencing, and playback. All sessions took place in the same open-topped plexiglass chamber (width, 28 cm; length, 28 cm; height, 20 cm), housed within a darkened, single-walled acoustic chamber (Industrial Acoustics, New York, NY) lined with anechoic foam [8,13,35]. No bedding, food, or water was provided. The chamber was cleaned with 70% isopropyl alcohol between sessions. Acoustic and video recording methods are described in subsequent sessions.

To record mating vocalizations, 16 animals (8 male-female pairs) were used in sessions that lasted for 30 minutes. A male mouse was introduced first into the test box, followed by a female mouse 5 min later. See details of acoustic video, and behavioral analyses below. To record vocalizations during restraint, six mice (4 male, 2 female) were briefly anesthetized with isoflurane and then placed in a restraint jacket as described previously [8,13]. Vocalizations were recorded for 30 minutes while the animal was suspended in the recording box.

See Figure 1 for the structure of playback experiments. All animals in these experiments (n = 36, 16 males, 20 females) underwent 90-min sessions on two consecutive days (Days 1 and 2) that provided both mating and restraint experiences (Fig. 1A). The order of experience sessions was presented in a counterbalanced pattern across subjects. Mating and restraint experiences were identical to those used in vocalization recording, except that the duration of the experiencing sessions was 90 mins. For the mating experience, mounting or attempted mounting was required for the animal to be included in the remainder of the experiment. However, we did not record detailed behaviors or track estrous stage during the mating experience session. An additional 36 mice (20 males, 16 females) were used as partners to the experimental animals in the mating experience but provided no vocalization or playback data.

**Figure 1.**
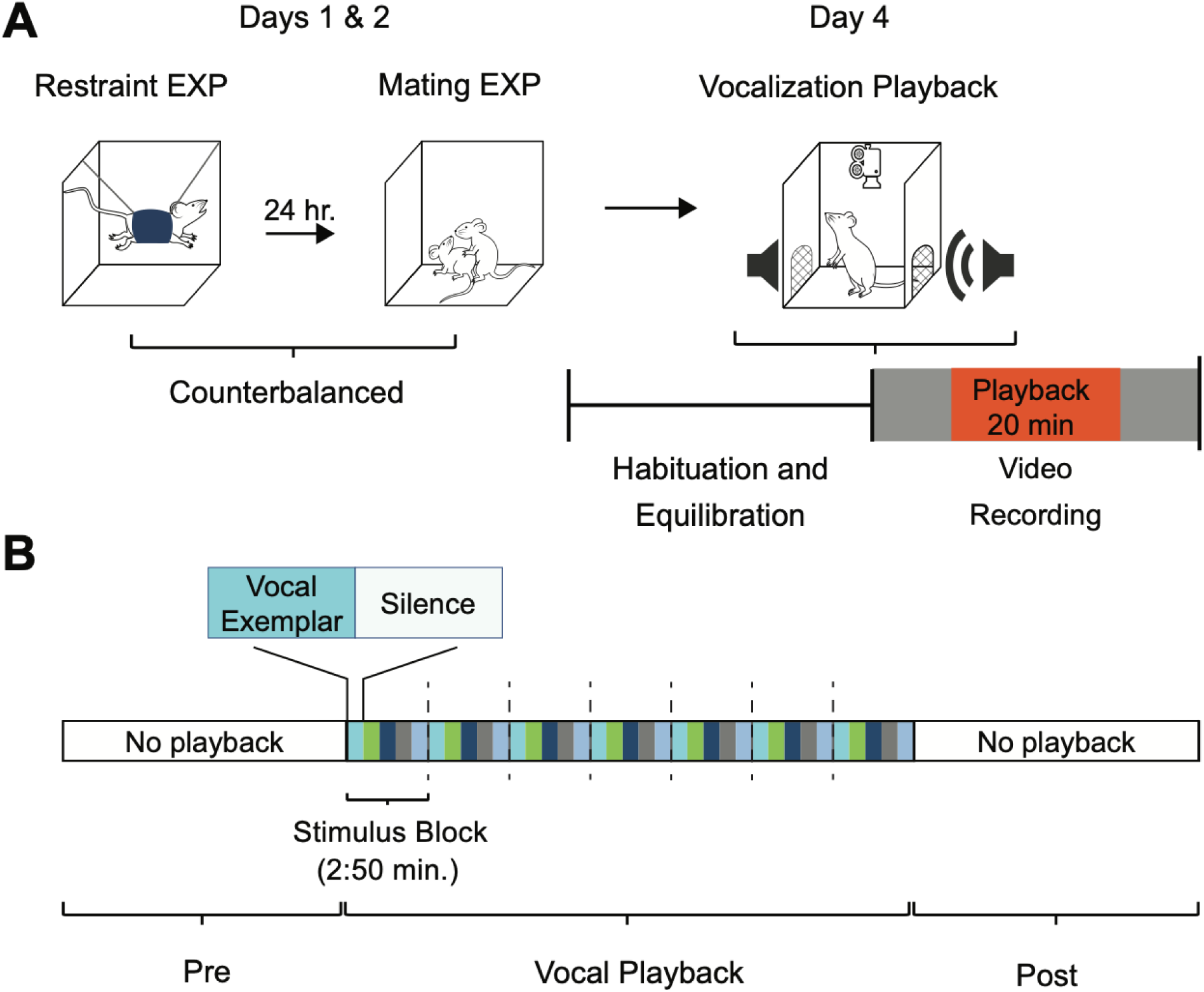
Temporal structures in playback experiment. **A.** Experimental sequence of experience and playback. On consecutive days, each animal experienced 90 min sessions of restraint and mating. The order of these sessions was counterbalanced across subjects. On the playback day, the subject was placed in the experimental chamber and allowed to habituate for three hours. Behavioral data was video-recorded for 40 mins: 10 mins before playback began, 20 mins during playback, and 10 mins after playback ended. **B.** Schematic illustration of detailed sequencing of vocal stimuli, shown here for mating playback. A 20-min period of vocal playback was formed by seven repeated stimulus blocks of 2:50 mins. The stimulus blocks were composed of five vocal exemplars (each represented by a different color) of variable length, with each exemplar followed by an equal duration of silence. In Figures 8-11, we report behavioral observations for blocks 1, 2, 3, 5, 6,and 7, omitting block 4. See Material and Methods for in-depth description of vocalization playback.

On Day 6, the day of the playback experiment, mice were randomly assigned to either restraint (8 males, 9 females) or mating (8 males, 11 females) vocal playback groups. Animals were placed within the experimental chamber and allowed to habituate for three hours, after which the 40 min behavioral observation period occurred.

Animal groups are shown in Table 1. Each animal in recordings experiments was used for one recording session and was not a subject in subsequent experiencing or playback experiments. Each animal in playback experiments was a subject for mating playback or restraint playback, but not both. However, all mice in playback experiments received both mating and restraint experiences. Additional mice were used to provide the opposite-sex partner in mating experiences, but these did not supply vocalization or playback data.

### 2.3 Acoustic methods

#### 2.3.1 Vocalization recording and analysis

Acoustic signals were recorded using ultrasonic condenser microphones (CM16/CMPA, Avisoft Bioacoustics, Berlin, Germany) connected to a multichannel amplifier and A/D converter (UltraSoundGate 416H, Avisoft Bioacoustics). The gain of each microphone was independently adjusted once per recording session to optimize the signal-to-noise ratio (SNR) while avoiding signal clipping. Acoustic signals were digitized at 500 kHz and 16-bit depth, monitored in real time with RECORDER software (Version 5.1, Avisoft Bioacoustics), and Fast Fourier Transformed (FFT) at a resolution of 512 Hz. A night vision camera (VideoSecu Infrared CCTV), centered 50 cm above the floor of the test box, recorded the behaviors synchronized with the vocal recordings (VideoBench software, DataWave Technologies, version 7).

Mating vocalizations were recorded using two ultrasonic microphones placed 30 cm above the floor of the recording box and 13 cm apart. See below for analysis of behaviors during vocal recordings. Since restraint vocalizations are usually emitted at lower intensity compared to mating vocalizations, the recording microphone was positioned 2-3 cm from the snout to obtain the best SNR.

Vocal recordings were analyzed offline using <u>Avisoft-SASLab Pro</u> (version 5.2.12, Avisoft Bioacoustics) with a hamming window, 1024 Hz FFT size, and an overlap percentage of 98.43. For every syllable the channel with the higher amplitude signal was extracted using a custom-written Python code and analyzed (https://github.com/GavazziDA/Wenstrup_Lab_Ghasemahmad_2023). Since automatic syllable tagging did not allow distinguishing some syllable types such as noisy calls and mid-frequency vocalizations (MFVs) from background noise, we manually tagged the start and end of each syllable, then examined spectrograms to measure several acoustic features and classify syllable types based on Grimsley and colleagues [13,18].

#### 2.3.2 Vocalization playback

Vocalization playback stimuli lasting 20 min were constructed from a sequence of seven repeating stimulus blocks lasting 2:50 min each (Fig. 1B). Each block was composed of a set of vocal sequence exemplars that alternated with an equal duration of background sound (“Silence”) associated with the preceding exemplar. The exemplars were recorded during mating interactions and restraint, and selected based on high SNR, correspondence with behavioral category by video analysis, and representation of the spectrotemporal features of vocalizations emitted during mating and restraint [13,36]. Mating stimulus blocks contained five exemplars of vocal sequences emitted during mating interactions, from five different male-female pairs. These exemplars ranged in duration from 15.0 – 43.6 s. Restraint stimulus blocks included seven vocal sequences, emitted by restrained male (n = 3) and female (n = 2) mice, with durations ranging from 5.7 – 42.3 s. Across exemplars, each stimulus block associated with both mating and restraint included different sets of vocal categories (Fig. 5).

Playback sequences, i.e., exemplars, were conditioned in Adobe Audition CC (2018), adjusted to a 65 dB SNR level, then normalized to 1V peak-to-peak for the highest amplitude syllable in the sequence. This maintained relative syllable emission amplitude in the sequence. For each sequence, an equal duration of background noise (i.e., no vocal or other detected sounds) from the same recording was added at the end of that sequence (Fig. 1B). A 5-ms ramp was added at the beginning and the end of the entire sequence to avoid acoustic artifacts. A MATLAB app (EqualizIR, Sharad Shanbhag; https://github.com/TytoLogy/EqualizIR) compensated and calibrated each vocal sequence for the frequency response of the speaker system. Vocal sequences were converted to analog signals at 500 KHz and 16-bit resolution using DataWave (DataWave SciWorks, Loveland, CO), anti-alias filtered (TDT FT6-2, fc=125KHz), amplified (HCA-800II, Parasound, San Francisco, CA), and sent to the speaker (LCY, K100, Ying Tai Audio Company, Hong Kong). Each sequence was presented at peak level equivalent to 85 dB SPL.

### 2.4 Behavioral methods

Behaviors during all sessions were recorded using a night vision camera (480TVL 3.6mm, VideoSecu), centered 50 cm above the floor of the test box, and SciWorks (DataWave, VideoBench version 7) for video acquisition and analysis.

#### 2.4.1 Male-female interactions for vocal recording

Interactions between male and female mice were video-recorded, then analyzed second-by-second. Among more general courtship interactions, we identified a set of behaviors that were mating-related, as described previously [11,37]. All vocal sequences selected as exemplars for playback of “mating” vocalizations were recorded in association with these male mating behaviors: head-sniffing, attempted mounting, or mounting (Table 2). Vocalizations during these behaviors included chevron, stepped, and complex USVs emitted with longer durations and higher repetition rates, and more LFH calls [11,36,38].

**Table 2:** Behaviors that distinguish courtship and mating. Based on previous work described in Section 2.4.1, these behaviors were used to distinguish courtship and mating behaviors and the vocalizations associated with each. Examples of these behaviors are shown in supporting video files 1-6.

| **Interaction Type** | **Behavior** | **Definition** |
| --- | --- | --- |
| **Courtship** | **Exploring** | Male and female explore the test box in the presence of the other sex. This behavior included rearing and sniffing corners and walls. |
|  | **Mutual sniffing** | Male and female sniffed each other's head, body or genital area simultaneously. |
|  | **Genital sniffing** | Male sniffs female's genital area. |
|  | **Female sniffing male's head** | Female sniffs male's head, snout, or face. |
| **Mating** | **Male sniffing female's head** | Male sniffs female's head, snout, or face. The behavior sometimes is followed by grooming of female's head. |
|  | **Attempt to mount/mount** | Male mouse mounts female followed by copulation behavior characterized by successive movement of his body while holding female with his forelimbs tightly for more than 2 seconds. If this behavior was interrupted by female's avoidance and lasted less than 2 seconds, mounting was considered unsuccessful (attempt to mount). Sniffing or approaching immediately before/after mounting or attempt to mount was considered as part of mounting behavior. |

#### 2.4.2 Experience and playback sessions

Behaviors were video-recorded for 40 mins, beginning 10 min before vocal playback (pre-stimulation, Pre), continuing for 20 min during playback (Vocal Playback), and including another 10 min after playback ended (post-stimulation, Post) (Fig. 6B). From the video recording, several behaviors (Table 3) were analyzed in 10-second intervals but reported in 2:50 min intervals associated with the blocks of vocal sequences. Since this did not result in an exact number of blocks, and as we aimed in capturing behavioral changes at same number of blocks at the beginning and end of playback window, we focused our statistical analysis and figures on the first and last three vocalization blocks (vocalization playback 1-3 and 4-7), excluding the middle block from the analysis.

**Table 3:**
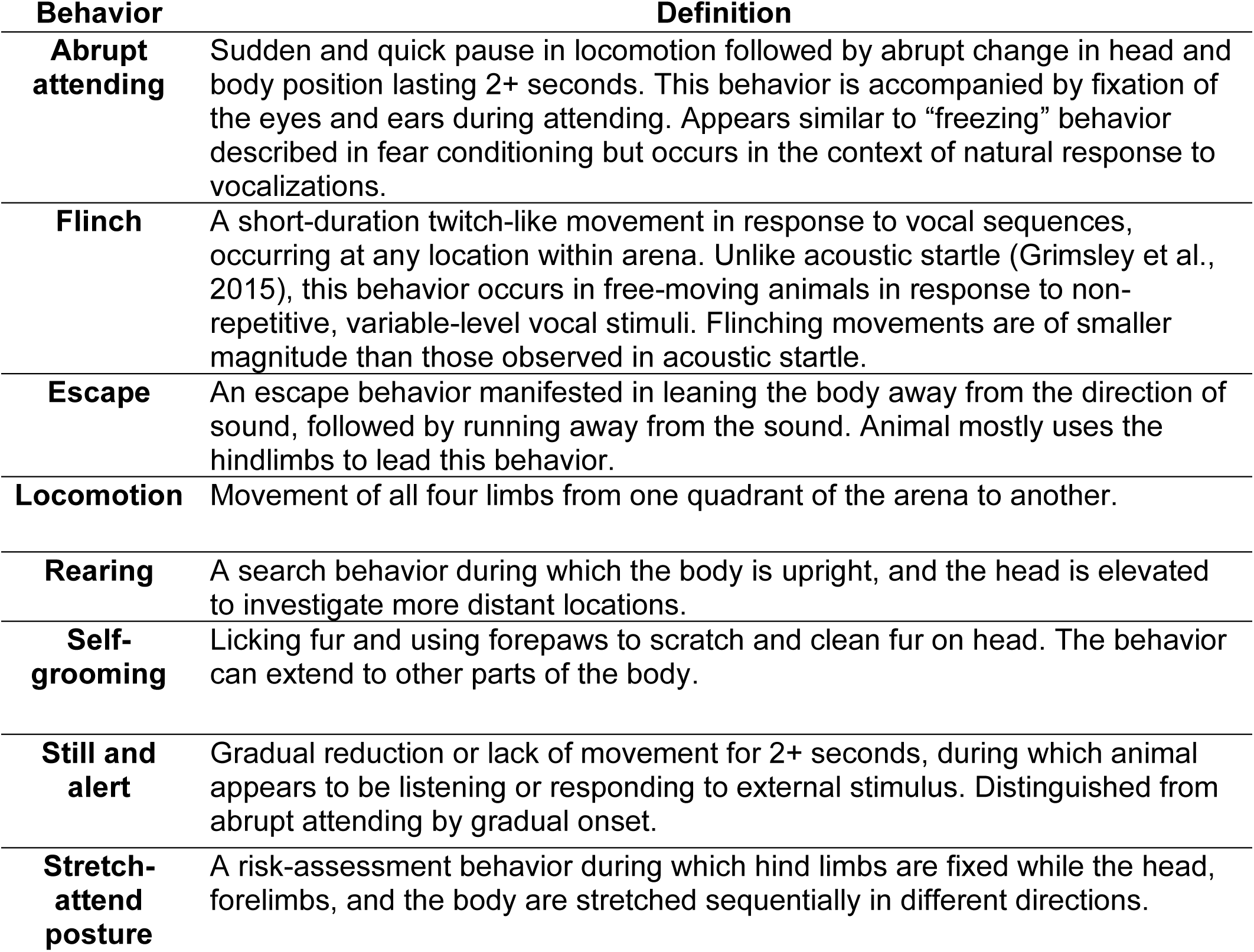
Classification of manually evaluated behaviors during vocalization playback. These behaviors are based on previous work as described in Section 2.4.2 and modified slightly from our previous study [8].

| **Behavior** | **Definition** |
| --- | --- |
| **Abrupt attending** | Sudden and quick pause in locomotion followed by abrupt change in head and body position lasting 2+ seconds. This behavior is accompanied by fixation of the eyes and ears during attending. Appears similar to “freezing” behavior described in fear conditioning but occurs in the context of natural response to vocalizations. |
| **Flinch** | A short-duration twitch-like movement in response to vocal sequences, occurring at any location within arena. Unlike acoustic startle (Grimsley et al., 2015), this behavior occurs in free-moving animals in response to non-repetitive, variable-level vocal stimuli. Flinching movements are of smaller magnitude than those observed in acoustic startle. |
| **Escape** | An escape behavior manifested in leaning the body away from the direction of sound, followed by running away from the sound. Animal mostly uses the hindlimbs to lead this behavior. |
| **Locomotion** | Movement of all four limbs from one quadrant of the arena to another. |
| **Rearing** | A search behavior during which the body is upright, and the head is elevated to investigate more distant locations. |
| **Self-grooming** | Licking fur and using forepaws to scratch and clean fur on head. The behavior can extend to other parts of the body. |
| **Still and alert** | Gradual reduction or lack of movement for 2+ seconds, during which animal appears to be listening or responding to external stimulus. Distinguished from abrupt attending by gradual onset. |
| **Stretch-attend posture** | A risk-assessment behavior during which hind limbs are fixed while the head, forelimbs, and the body are stretched sequentially in different directions. |

Behavioral analysis was based on previous descriptions [39–45] (Mouse Ethogram: https://mousebehavior.org) and consistent with our previous study [8]. The list of behaviors assessed was determined through pilot analysis and previous studies characterizing rodent defense and fear behaviors (Table 3). A behavior was only counted when its occurrence lasted two or more seconds, except for flinching, which could take place more quickly. Videos were further analyzed automatically using the video tracker within VideoBench (DataWave Technologies, version 7) for speed of locomotion, distance traveled, and time spent in the periphery and center of the test arena. All video analysis was performed blind to the sex of the animal and the context of vocalizations.

### 2.5 Data analysis

All statistical analyses were performed using SPSS (IBM, V. 26 and 27). To examine vocalizations in different stages of mating, we used a linear mixed model to analyze the changes of interval, duration, peak-to-peak amplitude and minimum frequency (dependent variables) of vocalizations based on mating interaction intensity, harmonicity, syllable types and the interaction of these (fixed effect). For behavioral analyses in playback experiments we used the General linear model with repeated measures for statistical comparisons of the behaviors. We used a full-factorial model of: intercept + sex + context + sex*context. Where Mauchly’s test indicated that the assumption of sphericity had been violated, degrees of freedom were corrected using Greenhouse Geisser estimates of sphericity.

To further clarify the differences observed between groups for every comparison in the GLM repeated measure, multivariate contrast was performed. All multiple comparisons were corrected using Bonferroni post hoc testing. 95% confidence intervals were used to compare values between timepoints [46].

For behavioral analyses, we tested the hypotheses that the number of behaviors was differently modulated by the vocal playback type (mating and restraint), by sex of the animals, or the interaction of these two. The GLM model for testing these hypotheses used time as a within-subject factor, with vocalization context, sex, and the interaction of these and time as the between-subject factors. Dependent (response) variables included the numbers of behaviors as previously described in the behavioral analysis section. In the analysis of stimulus-evoked behaviors, each animal’s pre-stimulus behavior served as a control. We did not use control groups presented with a non-social stimulus, since the valence or meaning of any artificial stimulus is uncertain [47,48].

To increase the analysis power, we focused the statistical analyses of the behaviors to the last 2:50 min block of pre-sound, first 2:50 block of vocal playback, and first 2:50 min of post playback. We chose a 2:50 analysis window based on the duration of a unique vocalization block and pre and post vocal playback windows were also chosen to match this window. Other time windows of behavioral observations, from other stimulus blocks, are shown in the figures but not used in statistical analysis.

## 3. RESULTS

The overall goal of these experiments was to examine the acoustic features of mouse social vocalizations associated with different, highly salient behavioral contexts—mating and restraint—and the behaviors in response to them. Vocalizations associated with restraint have been described previously [8,13] (see Figure 2A), and the development of restraint vocal sequences as stimuli was straightforward. Vocalizations associated with courtship and mating have also been extensively described [11,18,38] but this study required the development of vocalization stimuli typical of mating interactions. Experiment 1 describes the acoustic features of vocalizations emitted during mating, distinguished from less intense male-female courtship interactions. Some aspects of this analysis have been described previously [8]. Experiment 2 describes the behavioral responses to mating and restraint vocal sequences.

**Figure 2.**
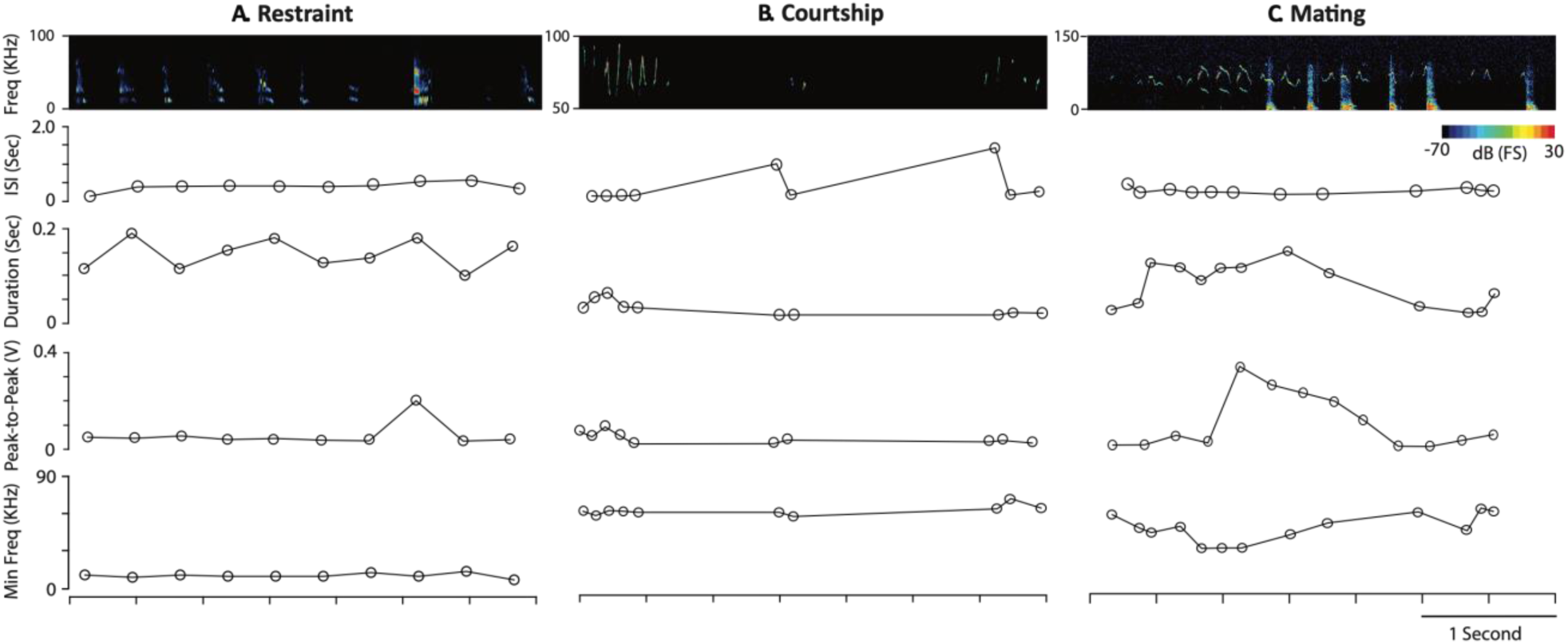
Vocalizations are modulated during different behavioral contexts. Each column displays short sequences of vocalizations and their acoustic features emitted by a single mouse in Restraint **(A)** or by CBA/CaJ male-female pairs in Courtship **(B)** and Mating **(C)**. Sound spectrograms (*top*) show syllables emitted during 3.5 s sequences in each behavior. Note different vertical axis scales for sonograms. Below the sonograms are four acoustic features of the vocal sequences: inter-syllable interval (ISI); duration (sec); sound level measured as peak-to-peak amplitude (volts); and minimum frequency (kHz). During Restraint, mice typically produce longer duration, lower frequency syllables such as the mid-frequency vocalization (MFV) with minimum frequencies below 18 kHz. Note that for Courtship (B) and Mating (C), graphs only show values for USVs. Compared to courtship, the higher intensity mating interaction is associated with more USV syllables and LFH calls, shorter ISI, longer duration, higher level, and lower minimum frequency.

### 3.1 Experiment 1: Acoustic features associated with intense mating interactions

We recorded behaviors and vocalizations (n = 13,293 syllables) of CBA/CaJ mice during 30-minute interactions between a male mouse and female mouse (n = 8 male-female pairs). We first categorized behaviors between the male-female pairs, separating these into categories associated with low intensity interactions (“courtship”) and high intensity interactions associated with attempted mounting (“mating”) (see Materials and Methods; [8,11,49]. Briefly, when female or male mouse engaged in mutual sniffing the vocalizations and behaviors were not intense (courtship), but when male sniffed female’s head, attempted to mount the female or had successful mounting, the behavior and the emitted vocalizations represented more intense interaction (mating). These behavioral classifications are shown in Table 2 and video examples are shown in Supporting Videos 1-6. During the 30-minute interactions, there were repeated bouts of mating (Fig. 3), but we saw no clear pattern throughout the 30-minute sessions or across mating pairs.

**Figure 3.**
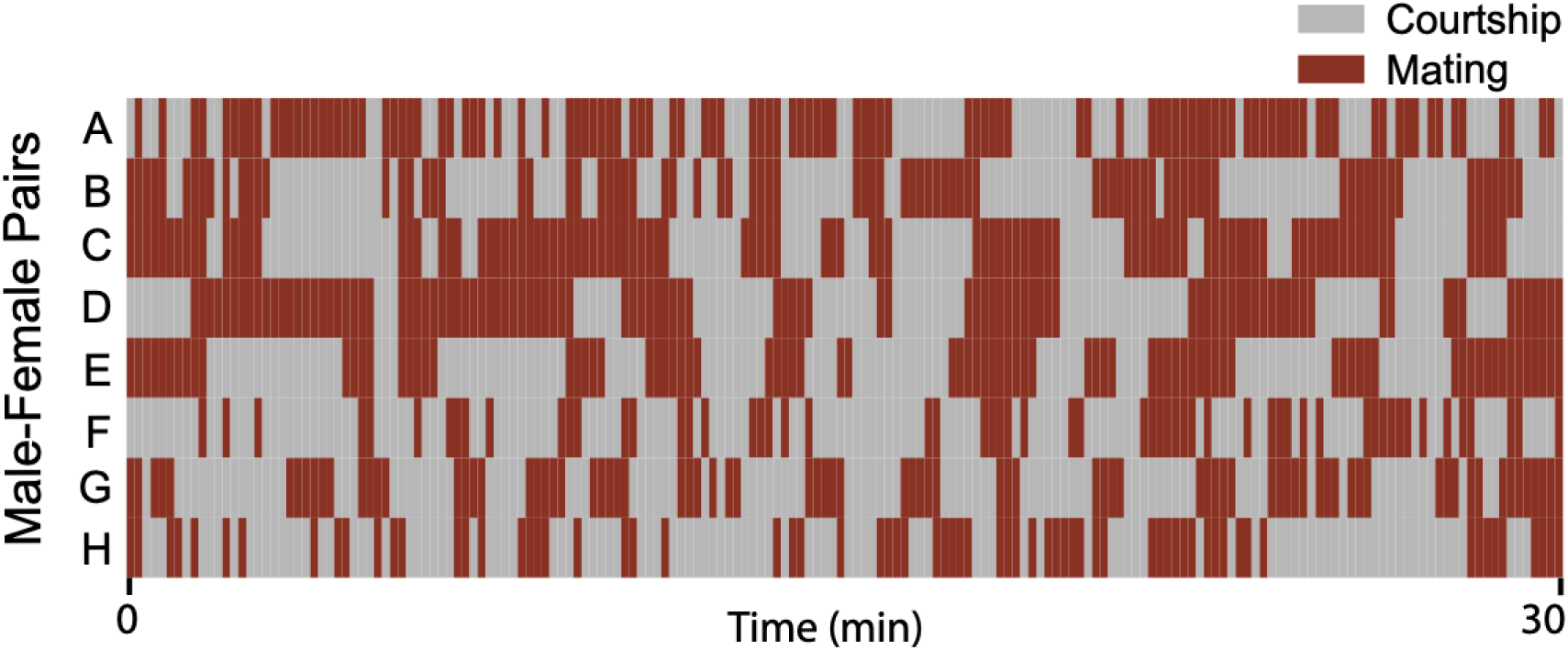
Sequences of courtship and mating behavior across male-female pairs. Each row represents one male-female pair for the 30-minute vocal recording. Times shown in red include any of the following behaviors associated with mating interactions (head-sniffing, attempt to mount, and mounting). Behaviors related to courtship (genital sniffing, mutual sniffing, exploring) are shown in grey.

We then analyzed the acoustic features of vocalizations emitted during these two behavioral categories. Example sequences and acoustic features are shown in Figures 2B and 2C. During courtship interactions, syllables consisting of mostly USVs were emitted in brief, relatively infrequent sequences, with long intervals between sequences (Fig. 2B). Vocal sequences emitted during mating consisted of both USVs, most likely emitted by the male, and low frequency harmonic (LFH) calls likely emitted by the female [11,19,38,50]. Compared to the courtship sequence, the mating sequence included more stepped and complex USVs, more fundamental harmonic components, lower minimum frequencies, longer syllable duration, and higher emitted sound levels (Fig. 2C). These general features of the vocalizations associated with these behavioral categories are documented in Figures 4-6. Both of these sequences differ substantially from those emitted during restraint (Fig. 2A).

**Figure 4.**
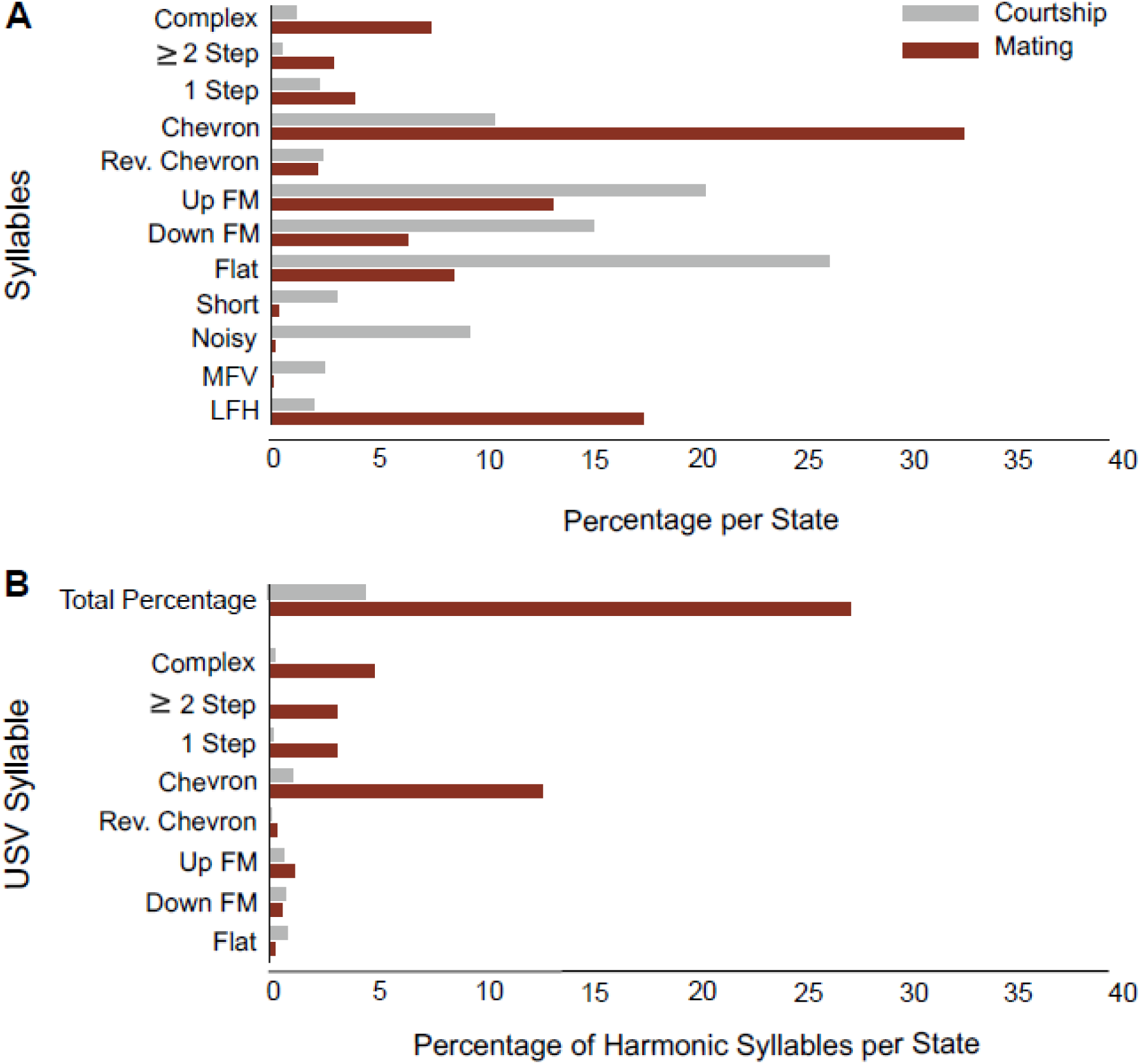
Syllable composition and harmonic structure change with male-female interaction intensity. **A.** Percentages of each syllable type recorded across courtship (gray) and mating (red) interactions. Note the higher percentage of complex, chevron, and stepped USV syllables and LFH calls emitted during mating. **B.** Frequency of occurrence of harmonics among USV syllables in courtship (gray) and mating (red) interactions. Note that many syllables shown in part A to be emitted preferentially during mating (i.e., complex, stepped, and chevron) have more harmonic components (syllables in courtship, n=6056; syllables in mating, n=7237 syllables).

Figure 4 compares syllable composition and harmonic content for vocalizations emitted during courtship and mating interactions. In this analysis, we include both USVs and non-USVs, whether or not the syllables overlapped, to represent the proportion of emitted syllables. During courtship interactions, the most common syllables were, in order: flat, up-FM, down-FM, chevron, and noisy (Fig. 4A). The most common syllables emitted during mating interactions other than female LFH calls, were mostly the same (chevron, up-FM, flat, and down-FM) but in a different order and very different proportions. Although syllable composition varied across the mating pairs (Supplemental Figure 1), there were several consistent patterns. Overall, mating USVs included more complex, stepped, and chevron syllables and fewer FM, flat, and short syllables, compared to courtship vocalizations. The mating syllables were much more likely to include at least one additional harmonic element, especially the chevron, complex, and stepped syllables (Fig. 4B). Syllable complexity thus increased during mating behavior, in agreement with previous studies [11,38,51]. To perform further analyses on spectral, temporal, and amplitude features of syllables, we removed from consideration all overlapping syllables, as occur when a male-emitted USV overlaps with a female-emitted LFH (see sonogram in Fig. 2C). The results of these analyses are shown in Figures 5 and 6.

**Figure 5.**
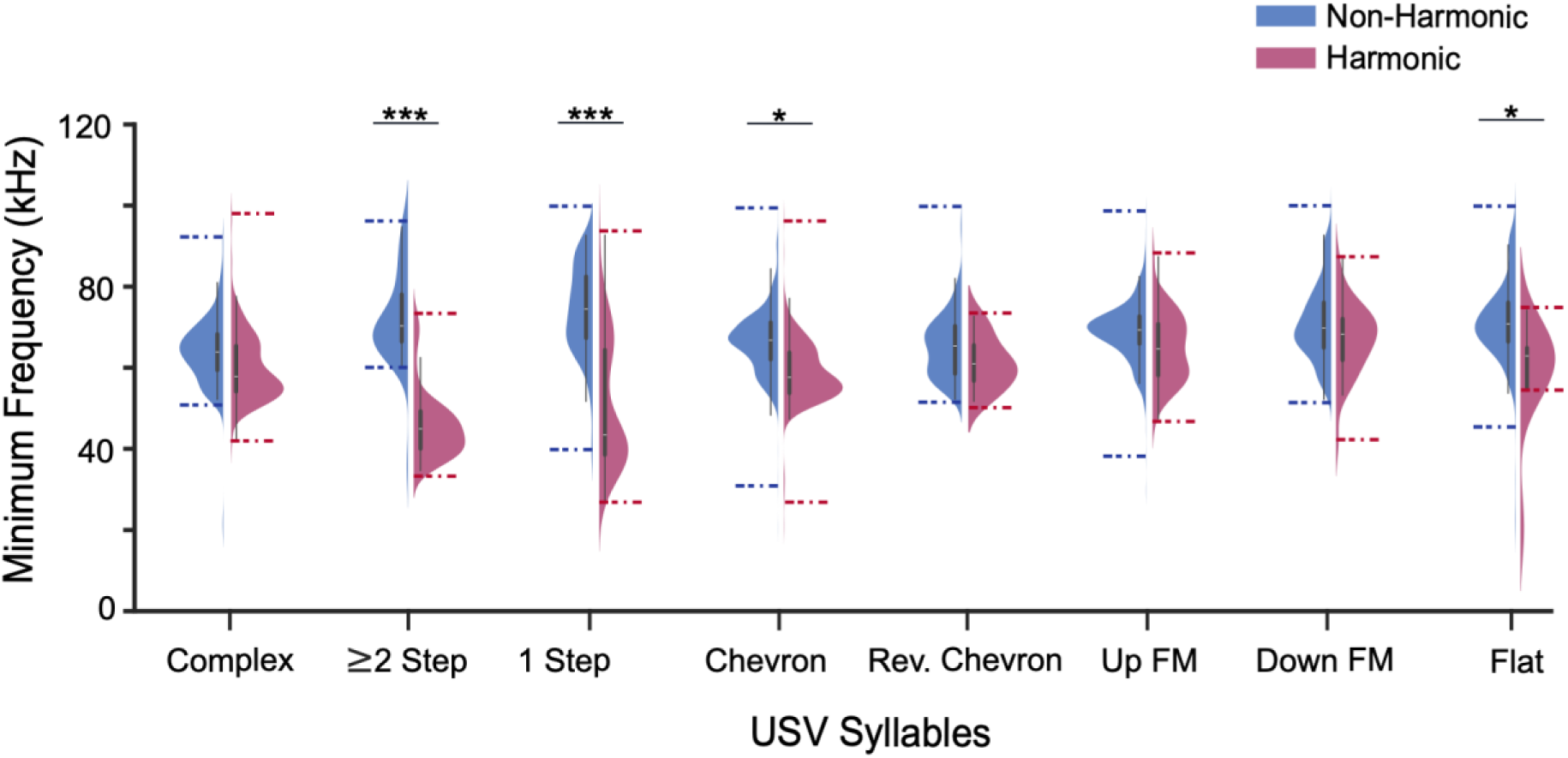
Minimum frequency of USVs decreases with increased intensity of mating interaction as a factor of harmonicity. Minimum frequency by syllable types based on the harmonicity (Significant three-way interaction of harmonicity, syllable type, and mating state on minimum frequency, n=4590 syllables, 9 USV syllable types, F_(7, 4582.2)_=4.0, *p<0.001*, Linear mixed model). Dashed lines represent the actual range of data for each syllable in harmonic and non-harmonic groups. Minimum frequency was significantly reduced for harmonic call in Flat (t =2.4, *p=0.02), Chevron (t=2.5, *p=0.01), 1 Step (t=10.63, \*\*\**p*<0.001), and ≥2 Step syllables (t=8.0, \*\**p*<0.001), but the effect in other syllables was not significant (e.g. UP FM: t =1.1, *p=0.3*, Down FM: t=1.4, *p*=0.16). Linear mixed model.

Previous work showed that the minimum frequency of syllables varies with the emotional state of the caller [11,13,38]. We found that, in male-female interactions, such context-related changes for USVs depended on syllable type and harmonicity (significant three-way interaction of mating state, syllable type, and harmonicity, n=4590 syllables, 9 USVs, F _(7, 4582.2)_ =4.0, *p<0.001*, Linear mixed model, Fig. 5). Thus, most of the syllables that were emitted more frequently during mating behavior (e.g., stepped and chevron syllables) and had multiple harmonics during mating also showed the most significant harmonicity-dependent reductions in minimum frequency (for instance one-stepped syllables t=10.63, *p<0.001*, chevron: t=2.5, *p=0.01*, or ≥2step syllables t=8.0, *p<0.001* Linear mixed model; Fig. 5). In contrast, other syllables that were emitted primarily in non-harmonic forms, such as up-FM and down-FM, showed little change in minimum frequency (e.g. up-FM: t = 1.1, *p=0.3*, down-FM: t=1.4, *p*=0.16, Linear mixed model; Fig. 5). The decreased minimum frequency that occurred with increased harmonicity appears to be due to the presence of the fundamental harmonic of these USVs (see Fig. 2C).

Temporal and amplitude features of vocalizations differed significantly with intensity of males-female interactions. Figure 6A shows syllable duration increased substantially during mating interaction in each male-female pair (n=8 mouse pairs, 10,717 syllables; F _(1,10715)_ =83.0, \*\*\**p<0.001,* Linear mixed model). Further, we observed a marked reduction in the inter-syllable interval (ISI) during mating interactions for all male-female pairs (significant main effect of state, n=8 mouse pairs and 10,189 syllable intervals: F _(1, 10180.2)_ =13.1, *** *p<0.001*, Linear mixed model; Fig. 6B). Sound amplitude also increased during mating; Figure 6C shows that peak-to-peak amplitude of all syllables increased across all mating pairs (n=8 mouse pairs, 10,313 syllables, F _(1,10307.0)_ =35.6, *\*\*\*p<0.001*, Linear mixed model). We excluded LFHs from amplitude analysis as these syllables are usually specific to the mating state and have significantly higher amplitude than other syllable types (Mean ± SD in volts: LFH (0.48±0.55), MFV (0.04±0.01), Noisy (0.06±0.04), Chevron (0.1±0.09)).

**Figure 6.**
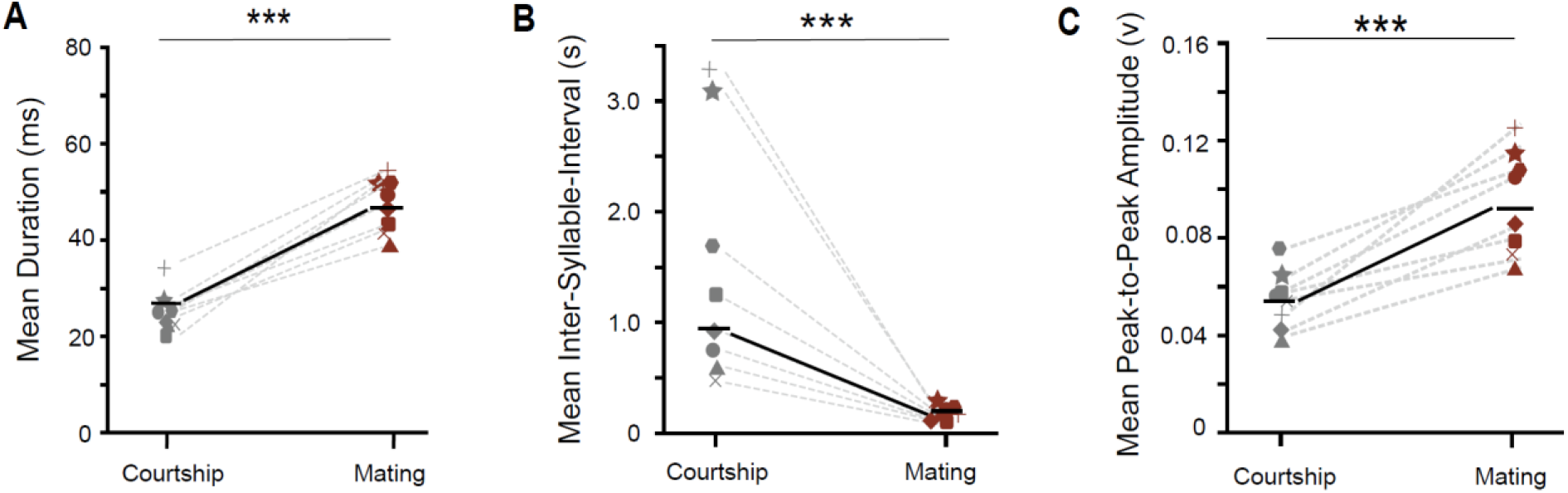
Temporal and amplitude features of syllables differ between courtship and mating interactions. Each symbol pair and connecting dashed line display average values for one mating pair in courtship and mating. Solid black line and symbols display average of all 8 mating pairs. **A**. Syllable duration increases with increased intensity of mating interaction (n=10,717 syllables; F(1,10715)=83.0, \*\*\**p*<0.001). **B**. Inter-Syllable-Interval (ISI) decreases with increased intensity of mating interactions (significant main effect of mating state, n=10,189 intervals: F(1, 10180.2)=13.1, *** *p*<0.001). **C**. Peak-to-peak amplitude of emitted syllables increases with mating intensity (n=10,313 syllables, F(1,10307.0)=35.6, \*\*\**p*<0.001). Linear mixed model.

For non-USV calls, there were clear differences between courtship and mating (Fig. 4A and Supplemental Fig. 1). LFH calls were much more common during mating, occurring during successful mounting by males and overlapping with USVs likely produced by males. By contrast, Noisy and MFV calls were much less frequent during mating than courtship.

Overall, our data suggest that during vocal communication in mice, the intensity of male-female interactions affects syllable composition (including USV, LFH, MFV, and noisy syllables), spectral features (harmonicity, minimum frequency), temporal features (e.g. interval and duration), and syllable intensity. Together with our previous study (Grimsley et al, 2016), these findings indicate that the change in state of vocalizing animals, represented in the intensity of social and sexual interactions, is reflected in the acoustic characteristics of their vocal signals.

Based on these findings and previous work by Grimsley et al. [13], we constructed vocal stimuli to be used in Experiment 2. Samples of these sequences are shown in Figure 7. Each spectrogram shows a 2-second sample of a longer sequence that contributed to the acoustic stimuli. Mating sequences (Fig. 7A) included many USVs with harmonics, steps, and complex structure, as well as LFH calls. Restraint vocalizations (Fig. 7B) include many USVs, MFVs, LFHs, and Noisy calls. These vocalizations were described previously in our related study on neuromodulator release [8].

**Figure 7.**
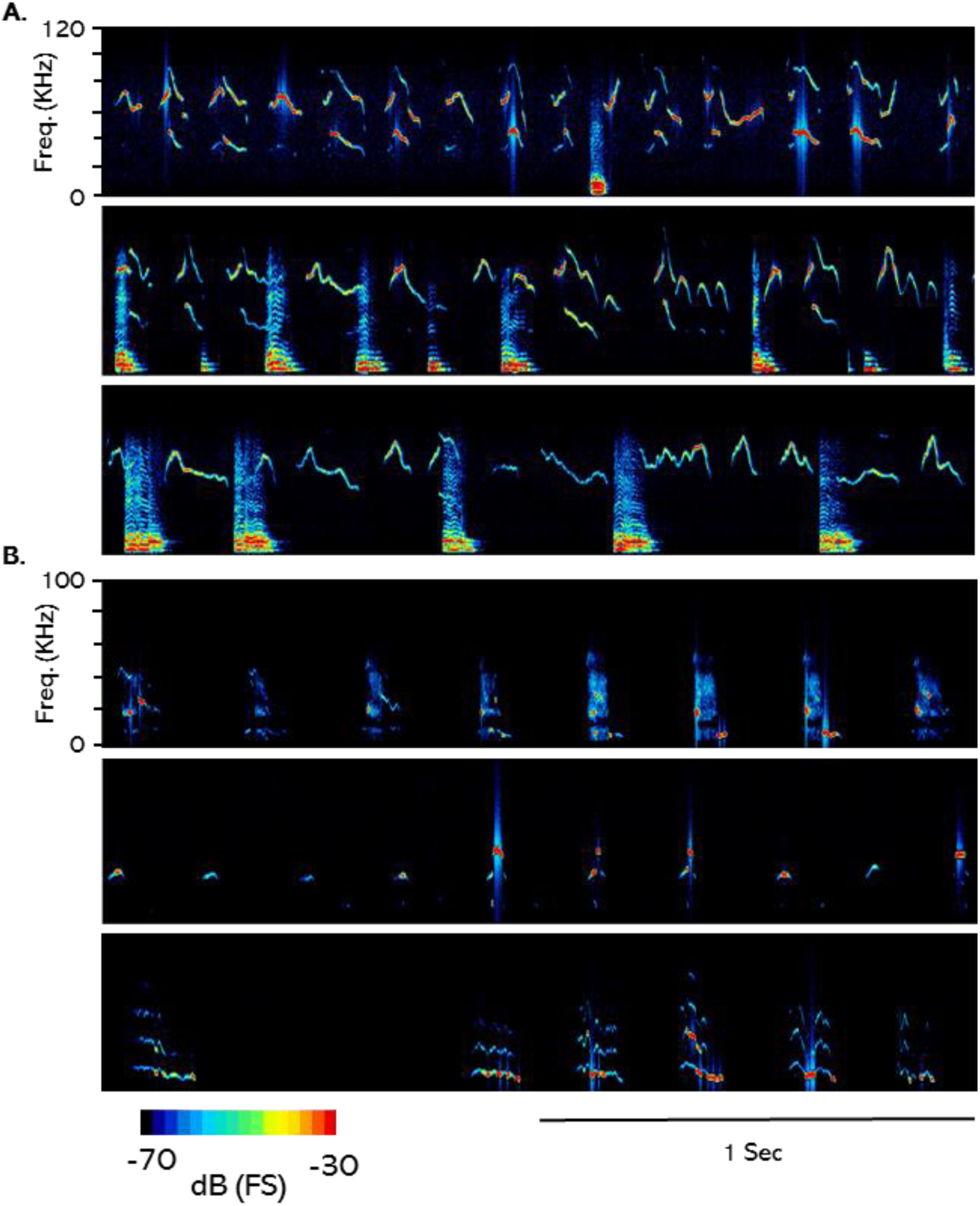
Spectrogram samples of vocal sequences used as stimuli in Experiment 2. **A.** Mating sequences. The durations of the full sequences were 5 s (top), 17 s (middle), and 7 s (bottom). **B.** Restraint sequences. The duration of the full sequences were 21 s (top), 18.2 s (middle), and 21 s (bottom).

### 3.2 Experiment 2: Emotional vocalizations result in behavioral changes in mice

In this experiment, we asked whether the emotionally charged vocalizations illustrated in Figure 7, representing a sender’s internal state in their syllable composition and spectrotemporal features, produced changes in the behavior of listening animals. We hypothesized that the high salience of our vocal stimuli would likely evoke some similar behaviors for both stimuli, while other behaviors would depend on the valence or behavioral context of the vocal stimuli. Further, since one context involves mating behavior, we hypothesized that there are sex-based differences in responses to these vocalizations. To test these hypotheses, we presented vocal stimulus blocks consisting of exemplars recorded either during the high-arousal stage of mating or during restraint, then monitored several behaviors of listening mice (n _female mating_= 9, n _male mating_= 8, n _female restraint_=11, n _male restraint_= 8). See Supporting Spreadsheet for behavioral data.

On the day of the playback experiment, mice acclimated to the experimental chamber for a period of three hours, after which we recorded their behaviors for 40 minutes during the following playback periods: Pre (10 mins) with no playback, Vocal Playback (20 mins) including playback of the entire group of mating or restraint sequences, and Post (10 mins) with no playback (Fig. 1B). Behavioral observations were collected continuously but are represented and analyzed in 2:50 min intervals equal to the duration of one mating or restraint stimulus block. In the following figures, we display behavioral measures throughout the 40 min recording period, in order to show the time course of behaviors before, during (six stimulus blocks), and after playback. However, our statistical analyses focus on three time points: 1) the last Pre interval, the first Vocal Playback interval, and the first Post interval.

In response to these two groups of contextual vocal sequences, mice displayed several patterns of behavioral responses. In one pattern, behavioral responses to playback were similar across both playback type and sex and were only observed during the playback periods. For instance, both male and female mice increased the number of attending behaviors (significant effect of sound: F _(1.37,43.76)_ =115.0, *p<0.001*; n=0.8) with vocalization playback regardless of the context (mating or restraint) associated with these stimuli (Vocal Playback vs Pre:, mean ± SD: female mating: +8.45± 1.4, restraint: +8.0± 4.8, male mating: +9.4.0± 3.0, restraint: +11.6± 5.1, Fig. 8A). Similarly, both male and female mice increased stretch-attend posture, a risk-assessment behavior associated in rodents with detection and analysis of possible threats [52,53]. This increase was significant in both sexes and contexts, except for males in response to mating playback calls (significant effect of sound: F _(1.07, 34.35)_ =44.4, *p*<0.001; n=0.6. see Fig. 8B). Overall females showed more increase in stretch-attend posture compared to males in mating playback), while this increase was equally observed in both male and female during restraint vocal playback (Vocal Playback vs Pre: female mating: +6.4 ±5.3, male mating: +2.75 ±3.5, female restraint: + 7.5 ±4.5, male restraint: + 6.25 ±5.0, Fig. 8B). Overall, both behaviors showed very low levels prior to the beginning of playback, with a sharp increase that was consistently observed during the first playback block. The behaviors were observed at lower levels in subsequent playback intervals and returned to baseline levels once playback ended (Fig. 8).

**Figure 8.**
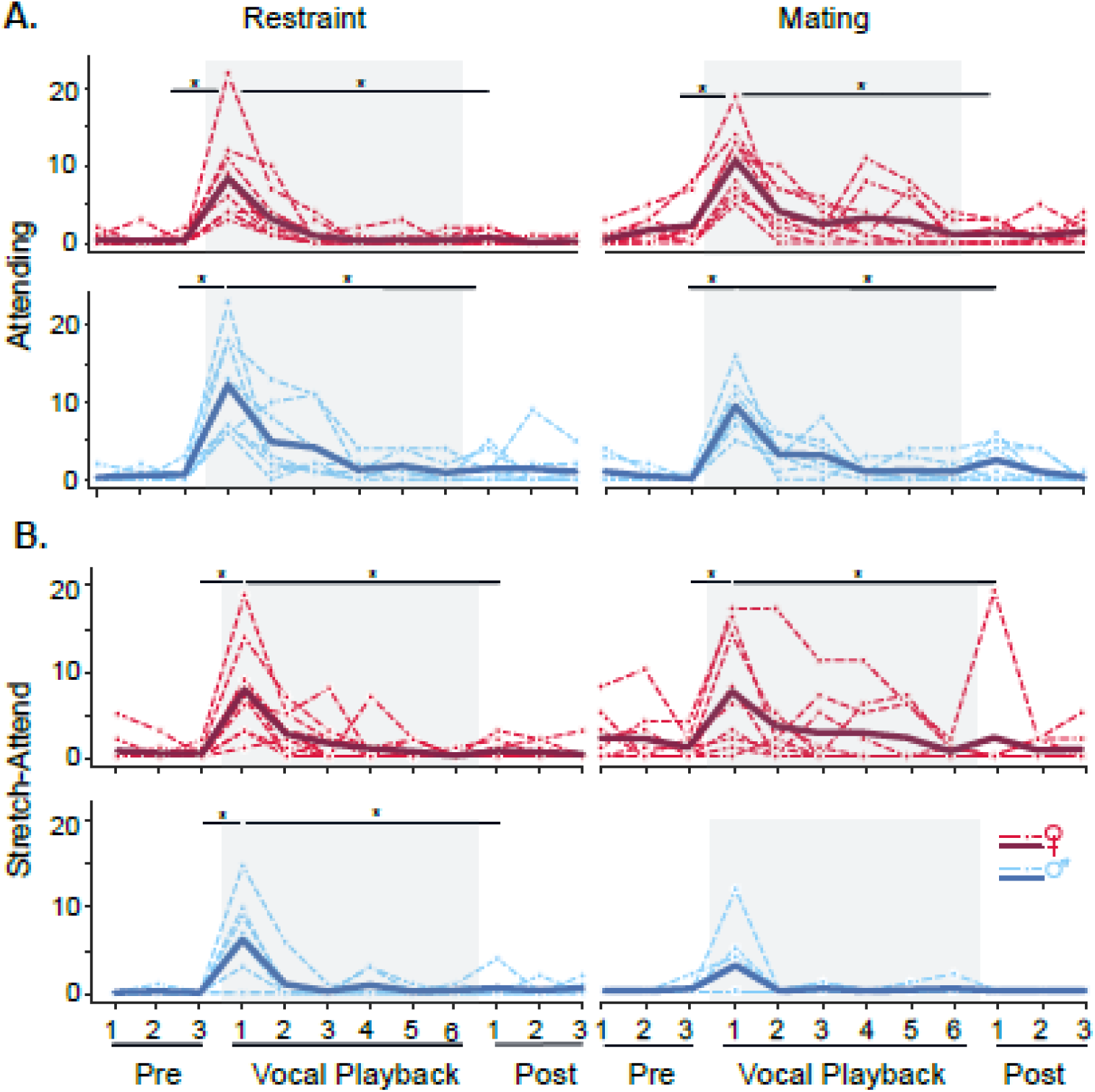
Playback of mating and restraint sequences evokes similar Attending and Stretch-Attend responses in male and female mice. For Figures 8-11, graphs display the number of behaviors during restraint (left) and mating (right) vocalization playback for estrus female (red) and male (blue) mice. Dashed lines represent individual animals; solid lines represent average number of the behavior for each group (n _female mating_=9, n _male mating_=8, n _female restraint_=11, n _male restraint_=8). **A.** The number of abrupt attending behaviors increased during both mating and restraint playback in both female and male mice (significant effect of vocal playback: F _(1.37,43.76)_ =115.0, *p<0.001*; n=0.8). **B**. Stretch-attend posture in female and male mice similarly increased during restraint and mating playback (significant effect of vocal playback: F _(1.07, 34.35)_ =44.4, *p*<0.001; n=0.6). Statistical tests based on repeated measures GLM with Bonferroni post hoc tests. *p<0.05, Time windows comparison: 95% confidence intervals.

Other behaviors seemed to be significantly shaped by the sex of the listening animal. For instance, females more than males showed reductions in self-grooming with start of vocal playback (significant main effect of sex: F (1,32) =4.28, p=0.04, Fig. 9A). Similarly, rearing decreased in females, accompanied by the increase of still-and-alert, a behavior signified by gradual reduction of activities (Fig. 9B, C). These behaviors displayed wider variation across individuals before, during, and after playback, suggesting that they were less closely tied to vocalization responses.

**Figure 9.**
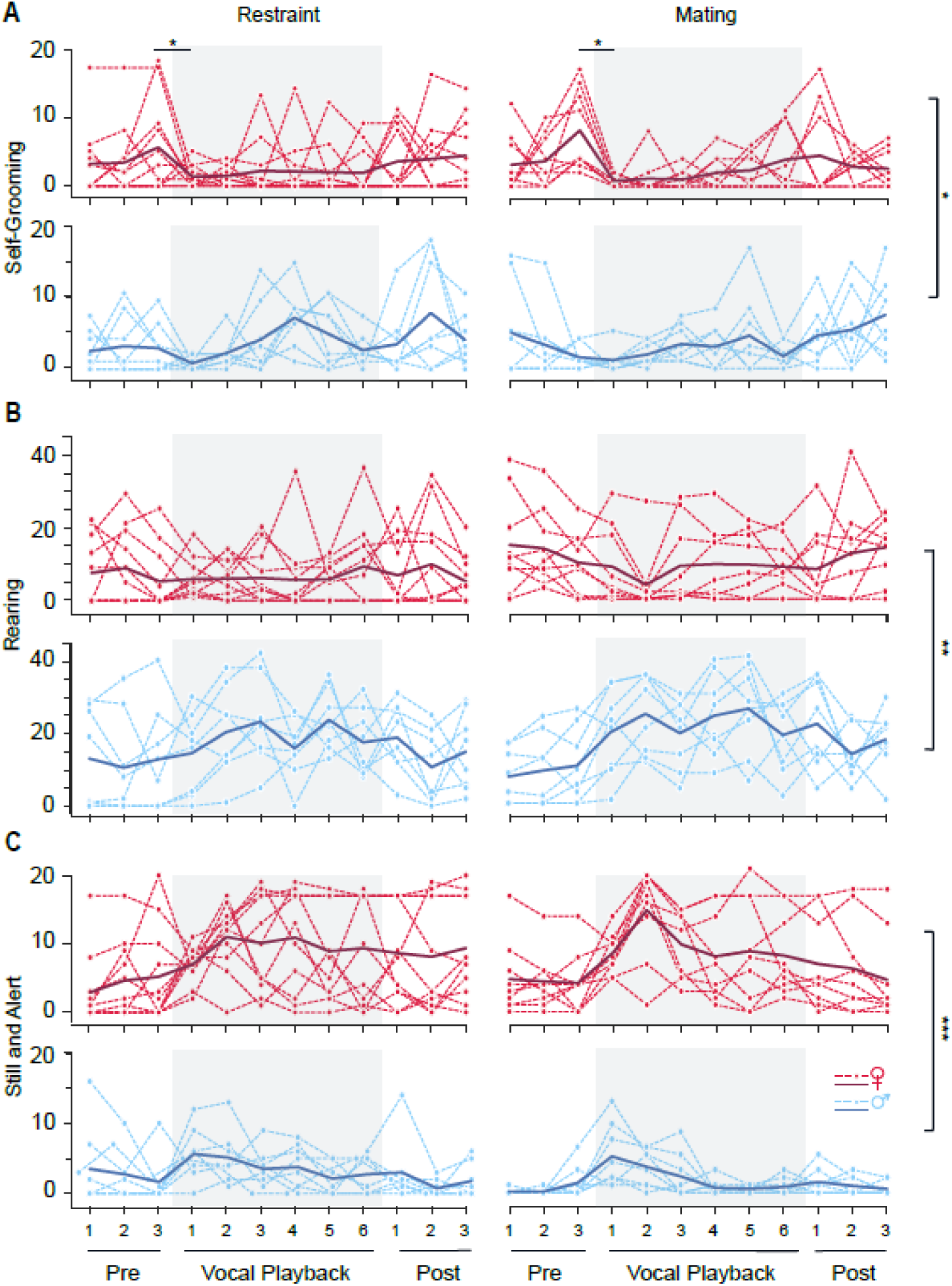
Playback of mating and restraint sequences evokes some behavioral responses that differ between male and female mice. See Figure 8 for protocol. **A.** Self-grooming behavior was reduced during playback in female mice more than males, regardless of vocalization type (significant main effect of sex: F _(1,32)_ =4.3, *P*=0.04; n=0.12). **B.** Rearing increased in males more than females regardless of vocalization type (F _(1,32)_ =10.6, *P*=0.003; n=0.25). **C.** Still and Alert behavior increased more in females than males during playback, regardless of vocalization type (F _(1,32)_ =16.7, *p*<0.001; n=0.34). *p<0.05, **p<0.01, ***p<0.001 (Bonferroni post hoc test).

Vocal playback also resulted in some behaviors that were shaped by the vocalization context. For instance, male and female mice flinched more during restraint playback, but not during mating playback (Fig. 10A) (significant interaction of time and context: F _(<u>1.02, 32.63</u>)_ =17.2, *p*<0. 001; n=0.<u>34</u>). This behavior was generally absent prior to vocal playback and followed a sound-dependent patterns in which the behavior diminished throughout playback duration. Locomotion behavior showed a significant increase during mating Vocal Playback (significant main effect of context: F _(<u>1, 32</u>)_ =8.0, *p*=0. 008; n=0.2) (Fig. 10B), with males generally showing more locomotion than females (Vocal Playback vs Pre: female mating: +1.2 ±1.5, male mating: +5.62 ±2, female restraint: +3.1 ± 4.4, male restraint: + 0.9 ± 0.1).

**Figure 10.**
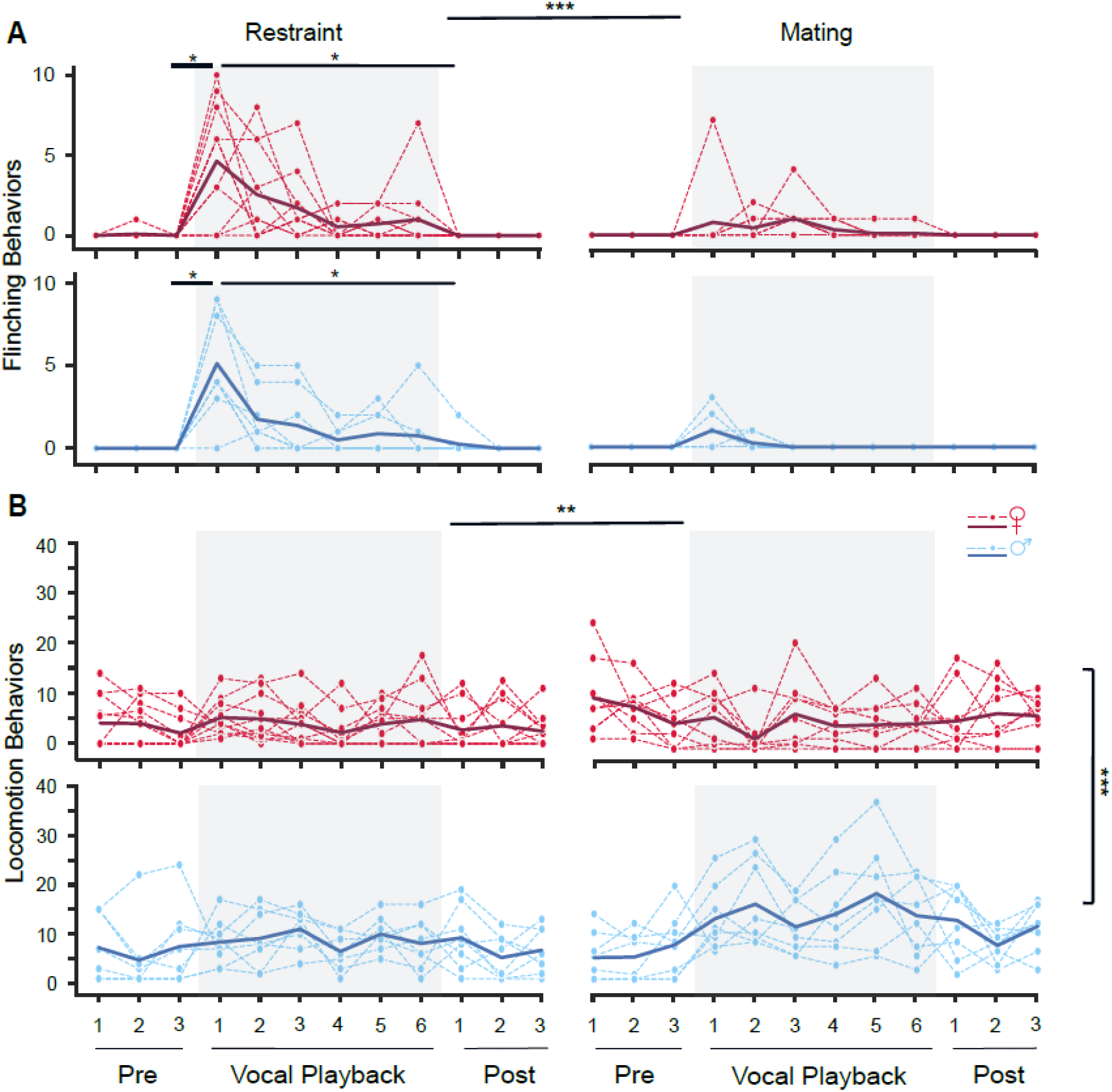
Playback of mating and restraint sequences evokes context-specific Flinching and Locomotion responses. See Figure 8 for protocol. **A.** Flinching behavior increased in both female and male mice during restraint but not in females during mating vocal playback (significant interaction of time and context: F _(1.02, 32.63)_ =17.2, *p*=<0.001; n=0.34). **B.** During mating playback, locomotion increased in male mice more than females (significant main effect of context: F_(1,32)_ =8.0, *p*=0.008; n=0.2). *p<0.05, **p<0.01, ***p<0.001. Time windows comparison: 95% confidence intervals.

Among the observed behaviors in these groups of animals, escape behavior showed a significant interaction between sex of the listener and the context of vocalizations, such that both mating and restraint vocalizations resulted in increased escape behavior in female mice, while males only exhibited such behavior during restraint vocalization playback (F _(1<u>.02, 32.6</u>)_ =5.0, *p*=0.03; n=0.13). Furthermore, this behavior in females showed more gradual decrease after the sound onset while in males was more abrupt after the first vocal playback block (Fig. 11).

**Figure 11.**
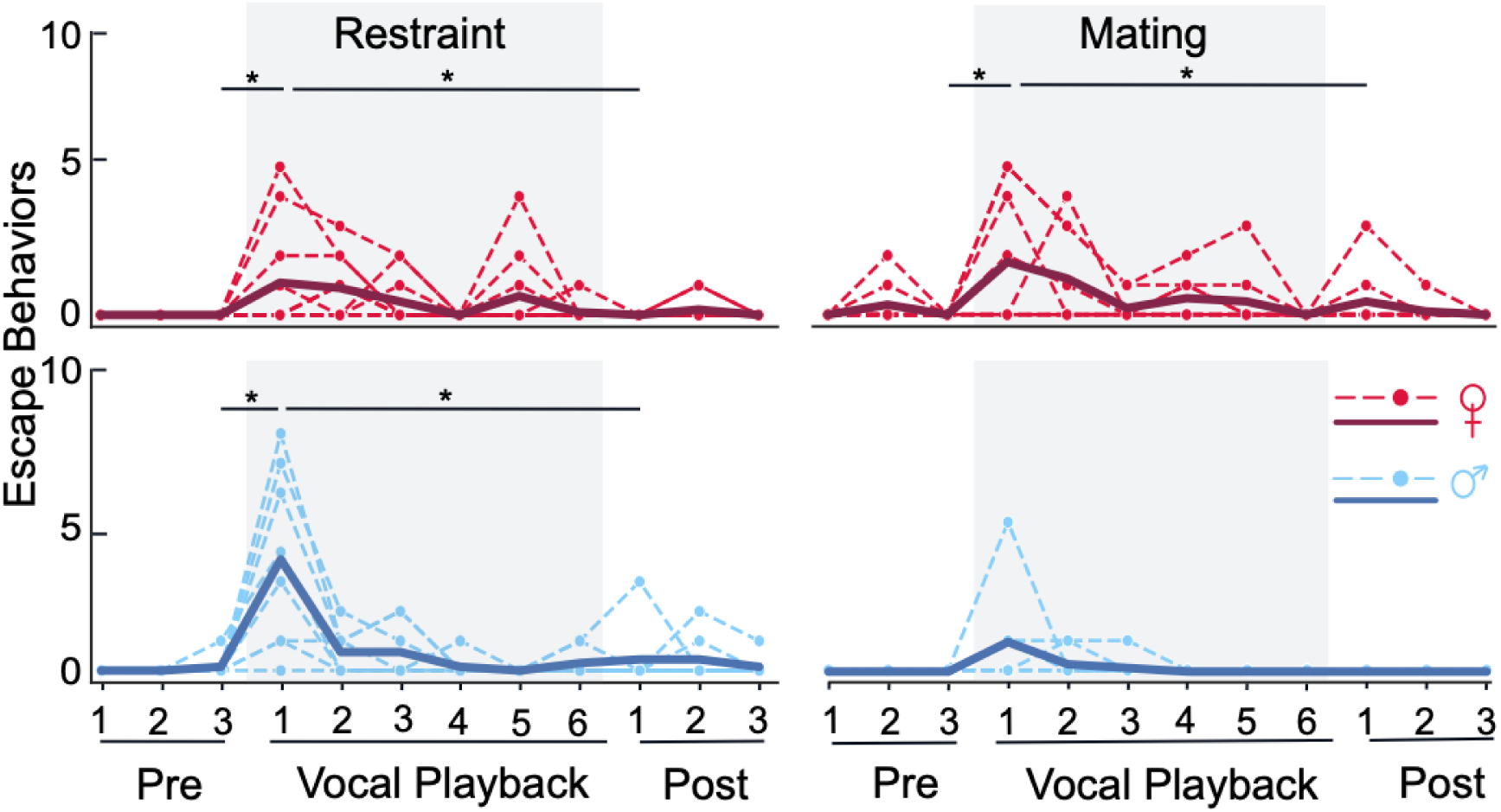
Escape behaviors differed by sex and vocal playback type. See Figure 8 for protocol. There was a significant interaction of time, sex, and context: F_(1.02, 32.6)_ =5.0, *p*= 0.03; n=0.13: repeated measures GLM, Bonferroni post hoc test, *p<0.05. Time windows comparison: 95% confidence intervals.

Overall, these behavioral findings show that salient vocalizations affect the behavior of listening mice. Further, the changes for some behaviors were independent of the context of vocalizations, but in other cases depended on the sex of listening animals, the context of vocalizations, or an interaction of these factors.

## 4. DISCUSSION

This study describes behavioral responses to emotional vocalizations produced by mice under very different behavioral contexts. The first experiment identified acoustic features associated with an appetitive social interaction, courtship and mating, that presumably reflect the internal state of vocalizing animals. We used multichannel audio and simultaneous video recordings to relate vocalizations to the intensity of male-female interactions. The behavioral observations provided the basis to distinguish vocal features associated with “courtship” and “mating”, then developed a series of vocal exemplars for use in the second (playback) experiment. With playback of highly salient, emotional vocalizations associated with mating and restraint, we observed a range of behavioral responses, some generalized to both types of stimuli, others sex-dependent, and others displaying context-by-sex interactions. Some of the behaviors were strongly linked to vocal playback while others appeared minimally linked. Clearly, vocalization-induced behavioral responses do not uniformly indicate an internal state change in the listening animal.

### 4.1 Vocalizations in courtship and mating

Vocalizations are valuable indicators of emotions or internal state that contribute to the affective component of social communication [54]. During vocal communications, both syllable type and spectrotemporal features are affected by a sender’s affective state. Change in fundamental frequency, duration, and rate of syllable emission have been reported in different species as they are affected by the internal state [3,11,13,55–58].

Motivated by the need to identify highly emotional vocal sequences associated with mating, we used previously described behaviors of male-female mouse pairs [11,49] to identify two levels of intensity of these interactions, “courtship” and “mating”. We then compared the acoustic features of a large number of vocalizations, over 13,000 syllables from 8 mouse pairs, that were produced during these two behavioral interactions. The mice emitted USVs and three lower frequency/broadband vocal categories. Our results show several changes in vocalizations as the intensity of the interaction increased. Large scale changes in vocalization sequences included increased repetition rate and sound level. There was also a change in overall syllable composition, marked by increased complexity of syllable types: more chevron, complex, and stepped calls, fewer FM, flat, and short calls). Especially for syllables that increased during the mating interactions, the duration and harmonic number increased while the minimum frequency decreased.

Our findings demonstrate that acoustic characteristics and syllable composition change with the intensity of male-female interactions in mice, presumably reflecting changes in the underlying internal state of the animals. These findings (change of syllable composition and their spectrotemporal features) are in general agreement with previous studies in other contexts [13] or similar context but different mouse models [11,59].

During mating interactions, chevrons were the dominant USV syllable, with increased proportion of syllables with frequency steps (40% vs 8%). These proportions differ from previous studies, only a few of which have identified chevron calls [13,18,38,60]. Instead, other studies report that syllables with frequency jumps are the main category emitted during the mating [11,16,17].

There are several possible explanations for this discrepancy. For instance, different mouse strains may utilize syllable types in different proportions [11,16,59,60]. Further, our recording techniques may have allowed us to detect chevron calls more sensitively. We used a multichannel recording setup for vocalization recordings. For every syllable, the channel with highest SNR was chosen to determine syllable type, rather than the single microphone recording used by others [16,59,60]. With the variations in head position during mating behaviors, a single microphone cannot represent the emitted spectrotemporal features of syllables as accurately, resulting in apparent discontinuities in the spectrograms. Thus, syllables will appear to include multiple elements or frequency steps rather than a continuous chevron shape. Finally, the necessity of removing overlapping syllables from further analysis [11,13,18] may have reduced the observation of chevron syllables, because these syllables are more likely to overlap with LFH calls during mating rather than courtship.

Although harmonicity changes have been reported in emotional vocalizations in various species [3,54,61–65], its covariation with other spectrotemporal characteristics of emotional vocalizations is rarely addressed [11,13]. The increase of syllable duration with harmonicity also occurs in vocalizations during aversive contexts in mice [13].

Our data show increased harmonicity of the major syllables (chevron and stepped calls) of mating, versus courtship. This change of harmonicity may result in the lowered fundamental frequency observed in our data during mating. Mating interactions also feature increased sound level of emitted syllables. Together, the higher sound levels and lower frequencies associated with harmonic syllables in mating will activate many more neurons in mouse Inferior colliculus [66], increasing the likelihood of responses in higher order regions such as the auditory cortex or amygdala.

Since longer duration syllables are the result of longer exhalations [67], and since USV production is tightly coordinated with sniffing behavior [68,69], changes in syllable duration and ISIs appear to be the result of a dramatic decrease in sniffing cycle [67,68,70]. Modulation of spectrotemporal characteristics of vocalizations that result from changes in vocal fold dynamics seem to be under tight control of sympathetic autonomic arousal [71,72]. They are most likely controlled by the central nucleus of amygdala in response to increased arousal [72].

It is noteworthy that the dramatic increase in emission of LFH calls by females occurred during mating sequences that included successful mounting by males. Although LFH calls are in some cases associated with female rejection of males, they also occur during successful mounting by males [8,19]. The LFH calls we recorded were mostly produced during successful mating sequences that were required for the male-female pair to be included in this vocalization analysis.

Overall, findings of this experiment help to identify the acoustic features that characterize different levels of appetitive male-female interactions leading to mating. We use these features to develop highly salient vocal stimuli with positive and negative affect that are used to study listeners’ behavioral responses to emotional vocalizations.

### 4.2 Behavioral responses to emotionally intense vocalizations associated with mating and restraint

Responses of mice and other animals to social vocalizations depend substantially on the multi-sensory and behavioral contextual features associated with these vocalizations. For instance, in mice, olfactory cues strongly influence behavioral responses to both USVs and LFH call, i.e., squeaks [11,38,47]. The types of social interactions (e.g., sex-dependency, aggression level) also influence the response to social vocalizations [23,73,74]. Because these multi-sensory and behavioral cues have important effects on acoustic communication, it has been difficult to assess how the vocalization itself shapes behaviors. This and our companion study [8] utilized playback of highly emotional vocalizations to study how the vocal stimuli alone shape listener’s response to vocal signals.

Our study controlled for or analyzed features of the listener that may affect behavioral responses. We ensured that all subjects had previous, well-defined experiences with the behaviors associated with the vocal stimuli since our parallel experiments showed that these experiences are critical to some neurochemical and behavioral responses [8]. To control for hormonal state, only females in estrus were included in the final analysis of the behavioral experiments. We further analyzed results by sex. Overall, the findings revealed some generalized responses that appear to result from enhanced attention and stimulus evaluation (e.g., increased attending–a freezing-like behaviors to stimuli of both contexts by both sexes), some sex-dependent responses (such as increased rearing in males to both vocal categories), and some differentially modulated by context (such as enhanced flinching or escape initiation responses in restraint playback in both sexes). Overall, salient mouse vocalizations that carry context-specific information in their composition and spectrotemporal attributes change the behaviors, and presumably the internal states, of listening mice. Further, the behaviors reveal both generalized responses to salient vocal signals and responses that are sex- or context-specific.

Findings of our current study agree with our previous work [8]. For example, in both studies, we observed an increase in attending in mice regardless of the type of vocal playback. The finer temporal analysis undertaken in the current study suggests that this Attending behavior is the result of the initial assessment of vocal stimuli, regardless of their type. This was revealed by fast decay of attending responses after the first playback period, in both sexes and with both playback types.

Consistent with our previous study, we further observed that the animal’s locomotive activity was substantially affected by sex of the listening animals during both behavioral contexts. In females, a response to vocal playback was to remain “still and alert”, reducing movements such as locomotion and rearing. In contrast, males generally were more active during vocal playback. This sex-based response to vocalization playback was only observed in playback of affective vocal sequences but not in playback of single syllables [31] or behavioral responses to other stimuli [75–77]. Further, females showed increased risk assessment behaviors (stretch-attend posture), suggesting cautious interpretation of vocal sequences. As females in our study were all in estrus, this behavioral response could be due to the amplified attention in estrus females, resulting from increased acetylcholine and potential interaction with dopamine across the estrous cycle [8].

Interestingly, in both of our studies, males tend to become more exploratory during mating vocal playbacks. In the current study, this is manifested in higher locomotion (exploring different parts of the arena) while in our previous study it appeared as increased rearing during mating vocal playback. Our previous study further suggests that such behavioral response is related to previous experience with this context, since males without mating experience did not show significant changes in motor responses during presentation of these vocal stimuli. As the mating vocal sequences used in our study included both the male and female vocalizations during mating stage, we cannot conclude which category of these vocalizations is responsible for triggering the observed behaviors. However, since the emotional context in our experiments are established through vocal sequence and syllable types rather than other sensory cues [4,32,47], and because these sequences have been associated with positive valence in vocalizing mice [11], we believe that the natural sequence of mating calls established a potential positive valence for the male mice, resulting in an increase in locomotion and exploratory behaviors throughout the playback period. Such active behaviors further represent males’ role during mating interactions, as male mice are overall more exploratory and active in examining the female and vocalizing to engage females in mating behavior [78]. Further, males did not exhibit avoidance or aversive behaviors such as escape initiation or flinching confirming the lack of negative cues in such vocalizations for male mice. Our neurochemical findings, indicating that dopamine release into the basolateral amygdala (BLA) in increased relative to other contexts, and acetylcholine (ACh) is decreased, further support an interpretation of positive valence for these sequences [8].

Both of our studies demonstrated significant interactions of sex and vocal type, in which vocal stimuli differentially modulated the behavior of male and female mice. In the present study, this response appeared as escape behavior, which males showed only during restraint playback while females showed regardless vocalization type. Our previous study, however, shows that differential influence of vocalization type on the sex of the listening animal can appear in other forms, such as flinching. There, we observed differential modulation of flinching behavior by the internal state of the animal (estrous phase) and vocalization type [8]. Such behaviors could be the result of vocalization type (e.g., responses to restraint vocalizations that were not influenced by restraint experience) or the result of the internal state of the listener (e.g., modulation by female’s estrus cycle). Overall, these studies demonstrate that both long terms features, i.e., sex and experience, and short-term factors such as hormonal state are critical to the listener’s response to these emotional vocalizations.

These results suggest further studies in two directions. First, the clear differences in vocal patterns during courtship and mating suggest tests to distinguish behavioral responses to playback of vocalizations emitted during these behaviors. Since in male-female interactions LFH calls are emitted mostly during actual or attempted mating, we expect that these calls may contribute to differences in behavioral responses to playback of mating and courtship vocal sequences, as other work has suggested [31,47,79,80]. These studies might distinguish the contributions of both LFH and the different USV patterns to behaviors during mating and courtship. A second direction might test what behavioral responses to acoustic sequences are common between vocal sequences and sequences of artificial sounds, such as broadband noise or tones. We did not use such sequences as control stimuli because there is evidence that they may not be emotionally “neutral” [47,48,81]. Nonetheless, additional studies might establish what behavioral responses to acoustic playback are the best indicators of internal responses to vocal differences, with potential effects of experience, sex, and hormonal state.

## Supporting information

Supplementary Figure 1 part 1

Supplementary Figure 1 part 2

Source data for Experiment II

Exploring video example

Mutual sniffing video example 1

Genital sniffing video example

Mutual sniffing video example 2

Head sniffing video example

Mounting video example

## CRediT AUTHORSHIP CONTRIBUTION STATEMENT

**Zahra Ghasemahmad:** Conceptualization, Methodology, Formal analysis, Investigation, Resources, Data curation, Writing – Original Draft, Writing – Review & Editing, Visualization, Supervision, Funding acquisition.

**Karthic Drishna Perumal:** Formal analysis, Writing – Review & Editing

**Bhavya Sharma:** Formal analysis, Writing – Review & Editing

**Rishita Panditi:** Formal analysis, Investigation, Writing – Review & Editing

**Jeffrey Wenstrup:** Conceptualization, Methodology, Resources, Data curation, Writing – Original Draft, Writing – Review & Editing, Visualization, Supervision, Project administration, Funding acquisition.

## ACKNOWLEDGMENTS

We are indebted to Sharad Shanbhag and Daniel Gavazzi for software used to condition vocal sequences and analyze audio and behavioral data, respectively, to Kristin Yeager of the Kent State University Statistical Consulting Center for statistical consulting, and to Sheila Fleming for advice on behavioral assessments. We thank Anthony Zampino, Austin Poth, Debin Lei, and Krish Nair for technical assistance. This work was supported by the National Institute on Deafness and Other Communication Disorders of the National Institutes of Health under award number R01DC00937-30 to J. Wenstrup and A. Galazyuk, and by an award from the Kent State University Graduate Student Senate to Z. Ghasemahmad.

## DATA AVAILABILITY

Software tools used in preparing sounds or analyzing sounds/behaviors are identified in the Materials and Methods and available on provided links to GitHub. Behavioral data is in the supporting data file. Vocalization data will be made available upon request to the authors.

## REFERENCES

[1] Y. Litvin, D.C. Blanchard, R.J. Blanchard, Vocalization as a social signal in defensive behavior, in: 2010: pp. 151–157. 10.1016/B978-0-12-374593-4.00015-2.

[2] S.M. Brudzynski, M. Silkstone, M. Komadoski, K. Scullion, S. Duffus, J. Burgdorf, R.A. Kroes, J.R. Moskal, J. Panksepp, Effects of intraaccumbens amphetamine on production of 50kHz vocalizations in three lines of selectively bred Long-Evans rats, Behavioural Brain Research 217 (2011) 32–40. 10.1016/j.bbr.2010.10.006.

[3] M.A. Gadziola, J.M.S. Grimsley, P.A. Faure, J.J. Wenstrup, Social Vocalizations of Big Brown Bats Vary with Behavioral Context, PLoS One 7 (2012). 10.1371/journal.pone.0044550.

[4] G. Ehret, Sound communication in house mice: Emotions in their voices and ears?, in: E. Altenmüller, S. Schmidt, E. Zimmermann (Eds.), Evolution of Emotional Communication: From Sounds in Nonhuman Mammals to Speech and Music in Man, First Edition, Oxford University Press, Oxford, 2013.

[5] S. Okabe, M. Nagasawa, T. Kihara, M. Kato, T. Harada, N. Koshida, K. Mogi, T. Kikusui, The Effects of Social Experience and Gonadal Hormones on Retrieving Behavior of Mice and their Responses to Pup Ultrasonic Vocalizations, Zoolog Sci 27 (2010) 790–795. 10.2108/zsj.27.790.

[6] S.M. Zala, D. Nicolakis, M.A. Marconi, A. Noll, T. Ruf, P. Balazs, D.J. Penn, Primed to vocalize: Wild-derived male house mice increase vocalization rate and diversity after a previous encounter with a female, PLoS One 15 (2020) e0242959. 10.1371/journal.pone.0242959.

[7] C.L. Petersen, S.E.D. Davis, B. Patel, L.M. Hurley, Social Experience Interacts with Serotonin to Affect Functional Connectivity in the Social Behavior Network following Playback of Social Vocalizations in Mice, ENeuro 8 (2021) ENEURO.0247-20.2021. 10.1523/ENEURO.0247-20.2021.

[8] Z. Ghasemahmad, A. Mrvelj, R. Panditi, B. Sharma, K.D. Perumal, J.J. Wenstrup, Emotional vocalizations alter behaviors and neurochemical release into the amygdala, Elife 12 (2024). 10.7554/eLife.88838.

[9] G.P. Lahvis, E. Alleva, M.L. Scattoni, Translating mouse vocalizations: Prosody and frequency modulation, Genes Brain Behav 10 (2011) 4–16. 10.1111/j.1601-183X.2010.00603.x.

[10] C.L. Giddens, K.W. Barron, J. Byrd-Craven, K.F. Clark, A.S. Winter, Vocal Indices of Stress: A Review, Journal of Voice 27 (2013) 390.e21–390.e29. 10.1016/j.jvoice.2012.12.010.

[11] S. Gaub, S.E. Fisher, G. Ehret, Ultrasonic vocalizations of adult male Foxp2-mutant mice: behavioral contexts of arousal and emotion, Genes Brain Behav 15 (2016) 243–259. 10.1111/gbb.12274.

[12] M. van Mersbergen, E. Lanza, Modulation of Relative Fundamental Frequency During Transient Emotional States, Journal of Voice 33 (2019) 894–899. 10.1016/j.jvoice.2018.07.020.

[13] J.M.S. Grimsley, S. Sheth, N. Vallabh, C.A. Grimsley, J. Bhattal, M. Latsko, A. Jasnow, J.J. Wenstrup, Contextual modulation of vocal behavior in mouse: Newly identified 12 kHz “Mid-frequency” vocalization emitted during restraint, Front Behav Neurosci 10 (2016). 10.3389/fnbeh.2016.00038.

[14] E. Lefebvre, S. Granon, F. Chauveau, Social context increases ultrasonic vocalizations during restraint in adult mice, Anim Cogn 23 (2020) 351–359. 10.1007/s10071-019-01338-2.

[15] D.T. Sangiamo, M.R. Warren, J.P. Neunuebel, Ultrasonic signals associated with different types of social behavior of mice, Nat Neurosci 23 (2020) 411–422. 10.1038/s41593-020-0584-z.

[16] T.E. Holy, Z. Guo, Ultrasonic songs of male mice, PLoS Biol 3 (2005) 1–10. 10.1371/journal.pbio.0030386.

[17] J. Chabout, A. Sarkar, D.B. Dunson, E.D. Jarvis, Male mice song syntax depends on social contexts and influences female preferences, Front Behav Neurosci 9 (2015). 10.3389/fnbeh.2015.00076.

[18] J.M.S. Grimsley, J.J.M. Monaghan, J.J. Wenstrup, Development of social vocalizations in mice, PLoS One 6 (2011). 10.1371/journal.pone.0017460.

[19] C.J. Finton, S.M. Keesom, K.E. Hood, L.M. Hurley, What’s in a squeak? Female vocal signals predict the sexual behaviour of male house mice during courtship, Anim Behav 126 (2017) 163–175. 10.1016/j.anbehav.2017.01.021.

[20] G. Ehret, Common rules of communication sound perception, in: J. Kanwal, G. Ehret (Eds.), Behavior and Neurodynamics for Auditory Communication, Cambridge University Press, New York, 2006: pp. 85–114.

[21] K.E. Radziwon, K.M. June, D.J. Stolzberg, M.A. Xu-Friedman, R.J. Salvi, M.L. Dent, Behaviorally measured audiograms and gap detection thresholds in CBA/CaJ mice, Journal of Comparative Physiology A 195 (2009) 961–969. 10.1007/s00359-009-0472-1.

[22] D.M. Green, T. Scolman, O.W. Guthrie, B. Pasch, A broad filter between call frequency and peripheral auditory sensitivity in northern grasshopper mice (Onychomys leucogaster), Journal of Comparative Physiology A 205 (2019) 481–489. 10.1007/s00359-019-01338-0.

[23] N.M. Rendon, S.M. Keesom, C. Amadi, L.M. Hurley, G.E. Demas, Vocalizations convey sex, seasonal phenotype, and aggression in a seasonal mammal, Physiol Behav 152 (2015) 143–150. 10.1016/j.physbeh.2015.09.014.

[24] S. Brudzynski, Handbook of mammalian vocalization: an integrative neuroscience approach., Academic Press, San Diego, 2009.

[25] M.L. Scattoni, J. Crawley, L. Ricceri, Ultrasonic vocalizations: A tool for behavioural phenotyping of mouse models of neurodevelopmental disorders, Neurosci Biobehav Rev 33 (2009) 508–515. 10.1016/j.neubiorev.2008.08.003.

[26] H.-X. Liu, O. Lopatina, C. Higashida, H. Fujimoto, S. Akther, A. Inzhutova, M. Liang, J. Zhong, T. Tsuji, T. Yoshihara, K. Sumi, M. Ishiyama, W.-J. Ma, M. Ozaki, S. Yagitani, S. Yokoyama, N. Mukaida, T. Sakurai, O. Hori, K. Yoshioka, A. Hirao, Y. Kato, K. Ishihara, I. Kato, H. Okamoto, S.M. Cherepanov, A.B. Salmina, H. Hirai, M. Asano, D.A. Brown, I. Nagano, H. Higashida, Displays of paternal mouse pup retrieval following communicative interaction with maternal mates, Nat Commun 4 (2013) 1346. 10.1038/ncomms2336.

[27] S. Valtcheva, R.C. Froemke, Neuromodulation of maternal circuits by oxytocin, Cell Tissue Res 375 (2019) 57–68. 10.1007/s00441-018-2883-1.

[28] S.M. Pomerantz, A.A. Nunez, N.J. Bean, Female Behavior is Affected by Male Ultrasonic Vocalizations in House Mice, Pergamon Press Ltd, 1983.

[29] C. V Portfors, D.J. Perkel, The role of ultrasonic vocalizations in mouse communication, Curr Opin Neurobiol 28 (2014) 115–120. 10.1016/j.conb.2014.07.002.

[30] K. Hammerschmidt, K. Radyushkin, H. Ehrenreich, J. Fischer, Female mice respond to male ultrasonic ‘songs’ with approach behaviour, Biol Lett 5 (2009) 589–592. 10.1098/rsbl.2009.0317.

[31] A.C. Niemczura, J.M. Grimsley, C. Kim, A. Alkhawaga, A. Poth, A. Carvalho, J.J. Wenstrup, Physiological and Behavioral Responses to Vocalization Playback in Mice, Front Behav Neurosci 14 (2020). 10.3389/fnbeh.2020.00155.

[32] K.L. Ronald, X. Zhang, M. V. Morrison, R. Miller, L.M. Hurley, Male mice adjust courtship behavior in response to female multimodal signals, PLoS One 15 (2020) e0229302. 10.1371/journal.pone.0229302.

[33] A.P.S. Dornellas, N.W. Burnham, K.L. Luhn, M. V. Petruzzi, T.E. Thiele, M. Navarro, Activation of locus coeruleus to rostromedial tegmental nucleus (RMTg) noradrenergic pathway blunts binge-like ethanol drinking and induces aversive responses in mice, Neuropharmacology 199 (2021). 10.1016/j.neuropharm.2021.108797.

[34] A.C. McLean, N. Valenzuela, S. Fai, S.A.L. Bennett, Performing vaginal lavage, crystal violet staining, and vaginal cytological evaluation for mouse estrous cycle staging identification, J Vis Exp (2012). 10.3791/4389.

[35] A.C. Niemczura, J.M. Grimsley, C. Kim, A. Alkhawaga, A. Poth, A. Carvalho, J.J. Wenstrup, Physiological and Behavioral Responses to Vocalization Playback in Mice, Front Behav Neurosci 14 (2020) 155. 10.3389/fnbeh.2020.00155.

[36] Z. Ghasemahmad, Behavioral and neuromodulatory responses to emotional vocalizations in mice, Kent State University, 2020. https://www.proquest.com/dissertations-theses/behavioral-neuromodulatory-responses-emotional/docview/2572567360/se-2?accountid=36418 (accessed March 19, 2023).

[37] J. Heckman, B. McGuinness, T. Celikel, B. Englitz, Determinants of the mouse ultrasonic vocal structure and repertoire, Neurosci Biobehav Rev 65 (2016) 313–325. 10.1016/j.neubiorev.2016.03.029.

[38] J.L. Hanson, L.M. Hurley, Female presence and estrous state influence mouse ultrasonic courtship vocalizations, PLoS One 7 (2012). 10.1371/journal.pone.0040782.

[39] V.P. Bakshi, A.E. Kelley, Feeding induced by opioid stimulation of the ventral striatum: role of opiate receptor subtypes, Journal of Pharmacology and Experimental Therapeutics 265 (1993) 1253–1260.

[40] V.P. Bakshi, A.E. Kelley, Sensitization and conditioning of feeding following multiple morphine microinjections into the nucleus accumbens, Brain Res 648 (1994) 342–346. 10.1016/0006-8993(94)91139-8.

[41] D.C. Blanchard, G. Griebel, R.J. Blanchard, The Mouse Defense Test Battery: pharmacological and behavioral assays for anxiety and panic, Eur J Pharmacol 463 (2003) 97–116. 10.1016/S0014-2999(03)01276-7.

[42] T. Füzesi, N. Daviu, J.I. Wamsteeker Cusulin, R.P. Bonin, J.S. Bains, Hypothalamic CRH neurons orchestrate complex behaviours after stress, Nat Commun 7 (2016) 11937. 10.1038/ncomms11937.

[43] C.A. Grimsley, R.J. Longenecker, M.J. Rosen, J.W. Young, J.M. Grimsley, A. V. Galazyuk, An improved approach to separating startle data from noise, J Neurosci Methods 253 (2015) 206–217. 10.1016/j.jneumeth.2015.07.001.

[44] K.R. Lezak, G. Missig, W.A. Carlezon Jr, Behavioral methods to study anxiety in rodents, Dialogues Clin Neurosci 19 (2017) 181–191. 10.31887/DCNS.2017.19.2/wcarlezon.

[45] P. Saibaba, G.D. Sales, G. Stodulski, J. Hau, Behaviour of rats in their home cages: daytime variations and effects of routine husbandry procedures analysed by time sampling techniques, Lab Anim 30 (1996) 13–21. 10.1258/002367796780744875.

[46] S.A. Julious, Using confidence intervals around individual means to assess statistical significance between two means, Pharm Stat 3 (2004) 217–222. 10.1002/pst.126.

[47] J.M.S. Grimsley, E.G. Hazlett, J.J. Wenstrup, Coding the meaning of sounds: Contextual modulation of auditory responses in the basolateral amygdala, Journal of Neuroscience 33 (2013) 17538–17548. 10.1523/JNEUROSCI.2205-13.2013.

[48] M.A. Gadziola, S.J. Shanbhag, J.J. Wenstrup, Two distinct representations of social vocalizations in the basolateral amygdala, J Neurophysiol 115 (2016) 868– 886. 10.1152/jn.00953.2015.

[49] J. Heckman, B. McGuinness, T. Celikel, B. Englitz, Determinants of the mouse ultrasonic vocal structure and repertoire, Neurosci Biobehav Rev 65 (2016) 313–325. 10.1016/j.neubiorev.2016.03.029.

[50] G. Oliveira-Stahl, S. Farboud, M.L. Sterling, J.J. Heckman, B. van Raalte, D. Lenferink, A. van der Stam, C.J.L.M. Smeets, S.E. Fisher, B. Englitz, High-precision spatial analysis of mouse courtship vocalization behavior reveals sex and strain differences, Sci Rep 13 (2023). 10.1038/s41598-023-31554-3.

[51] S.R. Egnor, K.M. Seagraves, The contribution of ultrasonic vocalizations to mouse courtship, Curr Opin Neurobiol 38 (2016) 1–5. 10.1016/j.conb.2015.12.009.

[52] D.C. Blanchard, G. Griebel, R. Pobbe, R.J. Blanchard, Risk assessment as an evolved threat detection and analysis process, Neurosci Biobehav Rev 35 (2011) 991–998. 10.1016/j.neubiorev.2010.10.016.

[53] T. Hager, R.F. Jansen, A.W. Pieneman, S.N. Manivannan, I. Golani, S. van der Sluis, A.B. Smit, M. Verhage, O. Stiedl, Display of individuality in avoidance behavior and risk assessment of inbred mice, Front Behav Neurosci 8 (2014). 10.3389/fnbeh.2014.00314.

[54] E. Zimmermann, L. Leliveld, S. Schenka, Toward the evolutionary roots of affective prosody in human acoustic communication: a comparative approach to mammalian voices., in: E. Altenmüller, S. Schmidt, E. Zimmermann (Eds.), Evolution of Emotional Communication: From Sounds in Nonhuman Mammals to Speech and Music in Man, Oxford University Press, Oxford, UK, 2013: pp. 117–132.

[55] B. Knutson, J. Burgdorf, J. Panksepp, Anticipation of play elicits high-frequency ultrasonic vocalizations in young rats., J Comp Psychol 112 (1998) 65–73. 10.1037/0735-7036.112.1.65.

[56] D. Rendall, M.J. Owren, Vocalizations as tools for influencing the affect and behavior of others, in: 2010: pp. 177–185. 10.1016/B978-0-12-374593-4.00018-8.

[57] A. Moles, F. Costantini, L. Garbugino, C. Zanettini, F.R. D’Amato, Ultrasonic vocalizations emitted during dyadic interactions in female mice: A possible index of sociability?, Behavioural Brain Research 182 (2007) 223–230. 10.1016/j.bbr.2007.01.020.

[58] J. Soltis,, Tracy E. Blowers, A. Savage, Measuring positive and negative affect in the voiced sounds of African elephants (*Loxodonta africana*), J Acoust Soc Am 129 (2011) 1059–1066. 10.1121/1.3531798.

[59] H. Wang, S. Liang, J. Burgdorf, J. Wess, J. Yeomans, Ultrasonic Vocalizations Induced by Sex and Amphetamine in M2, M4, M5 Muscarinic and D2 Dopamine Receptor Knockout Mice, PLoS One 3 (2008) e1893. 10.1371/journal.pone.0001893.

[60] Y.K. Matsumoto, K. Okanoya, Phase-specific vocalizations of male mice at the initial encounter during the courtship sequence, PLoS One 11 (2016). 10.1371/journal.pone.0147102.

[61] P.M. Narins, S.L. Smith, Clinal variation in anuran advertisement calls: basis for acoustic isolation?, Behav Ecol Sociobiol 19 (1986) 135–141. 10.1007/BF00299948.

[62] P.M. Narins, A.S. Feng, W. Lin, H.-U. Schnitzler, A. Denzinger, R.A. Suthers, C. Xu, Old World frog and bird vocalizations contain prominent ultrasonic harmonics, J Acoust Soc Am 115 (2004) 910–913. 10.1121/1.1636851.

[63] R.A. Suthers, P.M. Narins, W.-Y. Lin, H.-U. Schnitzler, A. Denzinger, C.-H. Xu, A.S. Feng, Voices of the dead: complex nonlinear vocal signals from the larynx of an ultrasonic frog, Journal of Experimental Biology 209 (2006) 4984–4993. 10.1242/jeb.02594.

[64] R. Márquez, Female Choice in the Midwife Toads (Alytes Obstetricans and a. Cisternasii), Behaviour 132 (1995) 151–161. 10.1163/156853995X00342.

[65] Benjamin E.F. Gourbal, M. Barthelemy, G. Petit, C. Gabrion, Spectrographic analysis of the ultrasonic vocalisations of adult male and female BALB/c mice, Naturwissenschaften 91 (2004). 10.1007/s00114-004-0543-7.

[66] M. Tehrani, S. Shanbhag, J.J. Huyck, R. Patel, D. Kazimierski, J.J. Wenstrup, The Mouse Inferior Colliculus Responds Preferentially to Non-Ultrasonic Vocalizations, ENeuro 11 (2024). 10.1523/ENEURO.0097-24.2024.

[67] G.A. Castellucci, D. Calbick, D. McCormick, The temporal organization of mouse ultrasonic vocalizations, PLoS One 13 (2018) e0199929. 10.1371/journal.pone.0199929.

[68] Y.B. Sirotin, M.E. Costa, D.A. Laplagne, Rodent ultrasonic vocalizations are bound to active sniffing behavior, Front Behav Neurosci 8 (2014). 10.3389/fnbeh.2014.00399.

[69] J.A. Alves, B.C. Boerner, D.A. Laplagne, Flexible Coupling of Respiration and Vocalizations with Locomotion and Head Movements in the Freely Behaving Rat, Neural Plast 2016 (2016) 1–16. 10.1155/2016/4065073.

[70] D.W. Wesson, J. V. Verhagen, M. Wachowiak, Why Sniff Fast? The Relationship Between Sniff Frequency, Odor Discrimination, and Receptor Neuron Activation in the Rat, J Neurophysiol 101 (2009) 1089–1102. 10.1152/jn.90981.2008.

[71] T. Johnstone, C.M. Van Reekum, T. Bänziger, K. Hird, K. Kirsner, K.R. Scherer, The effects of difficulty and gain versus loss on vocal physiology and acoustics, Psychophysiology 44 (2007) 827–837. 10.1111/j.1469-8986.2007.00552.x.

[72] A.M. Stewart, G.F. Lewis, K.J. Heilman, M.I. Davila, D.D. Coleman, S.A. Aylward, S.W. Porges, The covariation of acoustic features of infant cries and autonomic state, Physiol Behav 120 (2013) 203–210. 10.1016/j.physbeh.2013.07.003.

[73] E. Sasaki, Y. Tomita, K. Kanno, Sex differences in vocalizations to familiar or unfamiliar females in mice, R Soc Open Sci 7 (2020) 201529. 10.1098/rsos.201529.

[74] M. Premoli, V. Petroni, R. Bulthuis, S.A. Bonini, S. Pietropaolo, Ultrasonic Vocalizations in Adult C57BL/6J Mice: The Role of Sex Differences and Repeated Testing, Front Behav Neurosci 16 (2022). 10.3389/fnbeh.2022.883353.

[75] M. Kent, M. Bardi, A. Hazelgrove, K. Sewell, E. Kirk, B. Thompson, K. Trexler, B. Terhune-Cotter, K. Lambert, Profiling coping strategies in male and female rats: Potential neurobehavioral markers of increased resilience to depressive symptoms, Horm Behav 95 (2017) 33–43. 10.1016/j.yhbeh.2017.07.011.

[76] I.R. Bishnoi, K. Ossenkopp, M. Kavaliers, Sex and age differences in locomotor and anxiety-like behaviors in rats: From adolescence to adulthood, Dev Psychobiol 63 (2021) 496–511. 10.1002/dev.22037.

[77] M.F. Herselman, L. Lin, S. Luo, A. Yamanaka, X.-F. Zhou, L. Bobrovskaya, Sex-Dependent Effects of Chronic Restraint Stress on Mood-Related Behaviours and Neurochemistry in Mice, Int J Mol Sci 24 (2023) 10353. 10.3390/ijms241210353.

[78] J.A. Johansen, L.G. Clemens, A.A. Nunez, Characterization of copulatory behavior in female mice: Evidence for paced mating, Physiol Behav 95 (2008) 425–429. 10.1016/j.physbeh.2008.07.004.

[79] Y. Chen, Q.Q. Su, Q.S. Liu, Effects of quinestrol on the vocal behavior of mice during courtship interactions, Physiol Behav 173 (2017) 216–222. 10.1016/j.physbeh.2017.02.017.

[80] K.E. Hood, E. Long, E. Navarro, L.M. Hurley, Playback of broadband vocalizations of female mice suppresses male ultrasonic calls, PLoS One 18 (2023). 10.1371/journal.pone.0273742.

[81] A.J. Parsana, N. Li, T.H. Brown, Positive and negative ultrasonic social signals elicit opposing firing patterns in rat amygdala, Behavioural Brain Research 226 (2012) 77–86. 10.1016/j.bbr.2011.08.040.

