## Supplementary figures and images for "Acoustic features of and behavioral responses to emotionally intense mouse vocalizations"

### Supplementary Figure 1 part 1

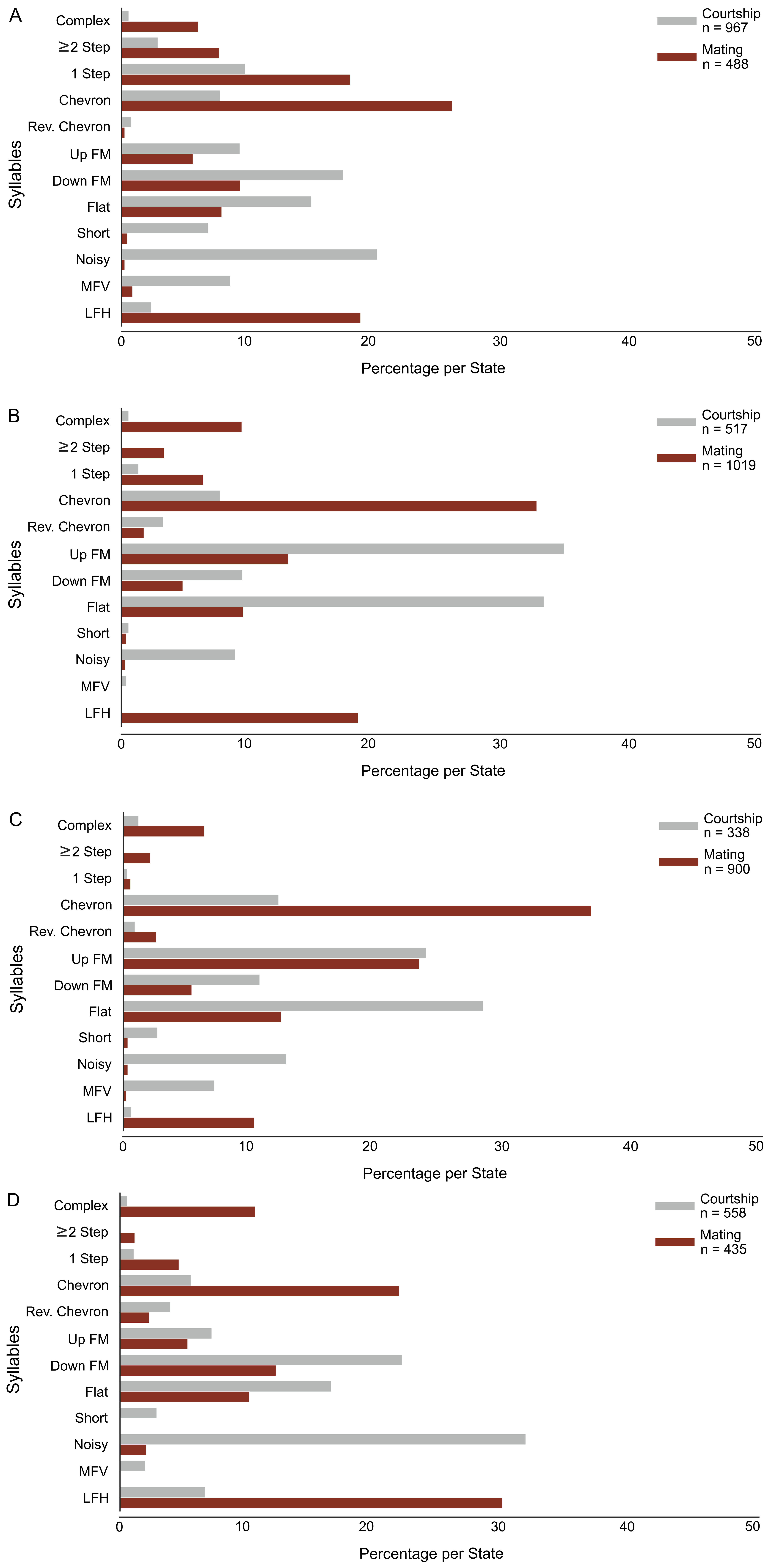

### Supplementary Figure 1 part 2

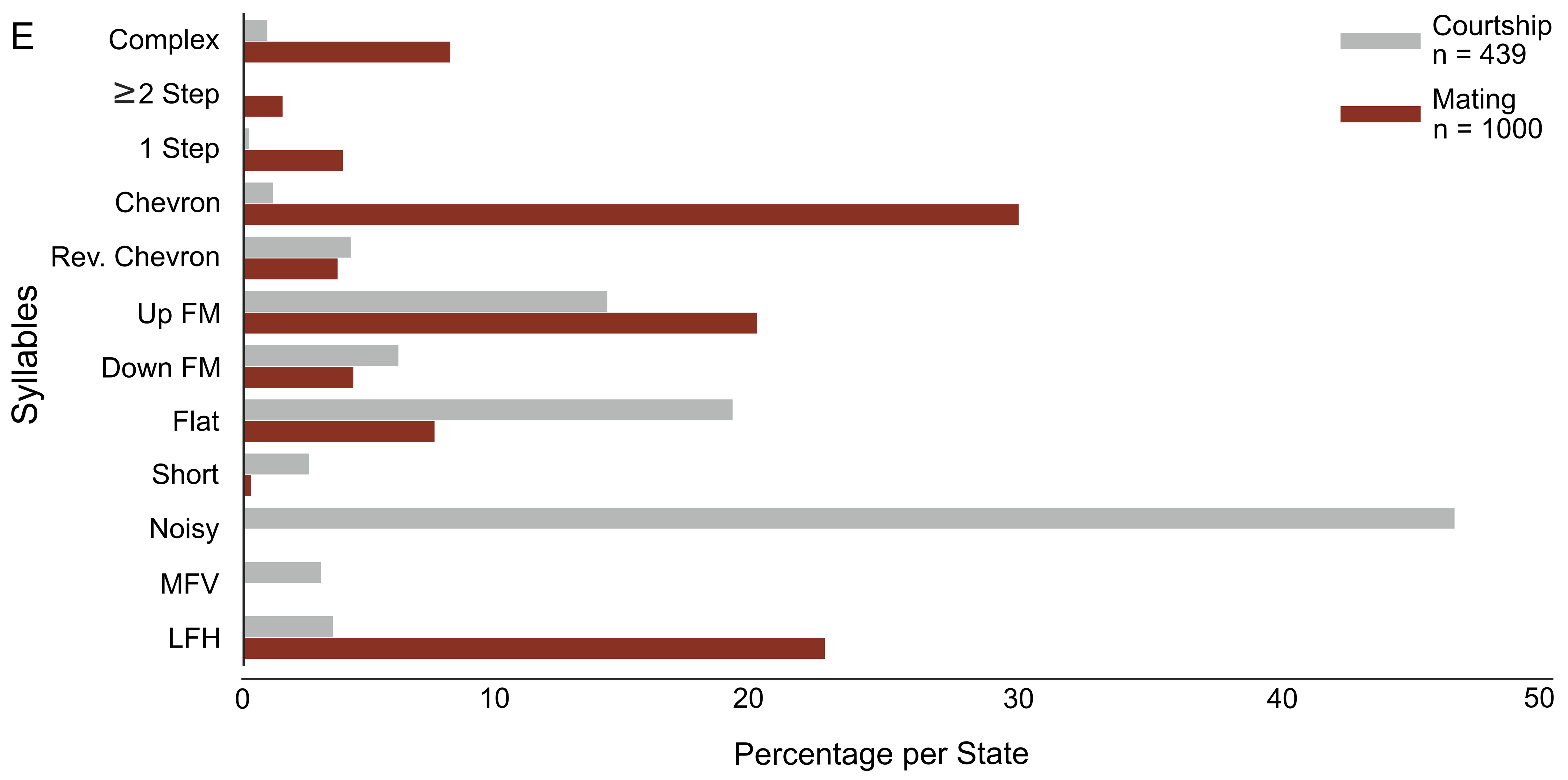

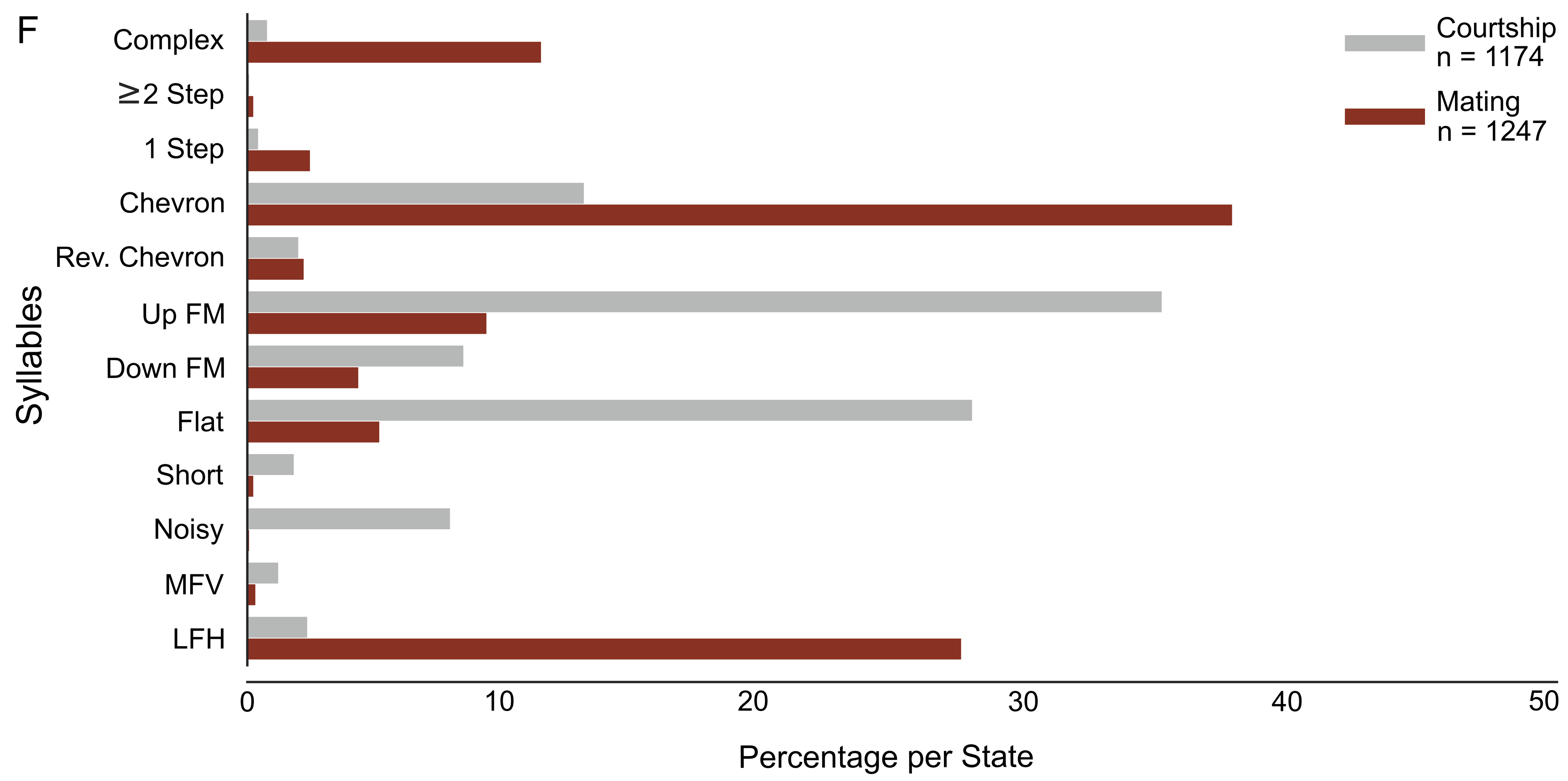

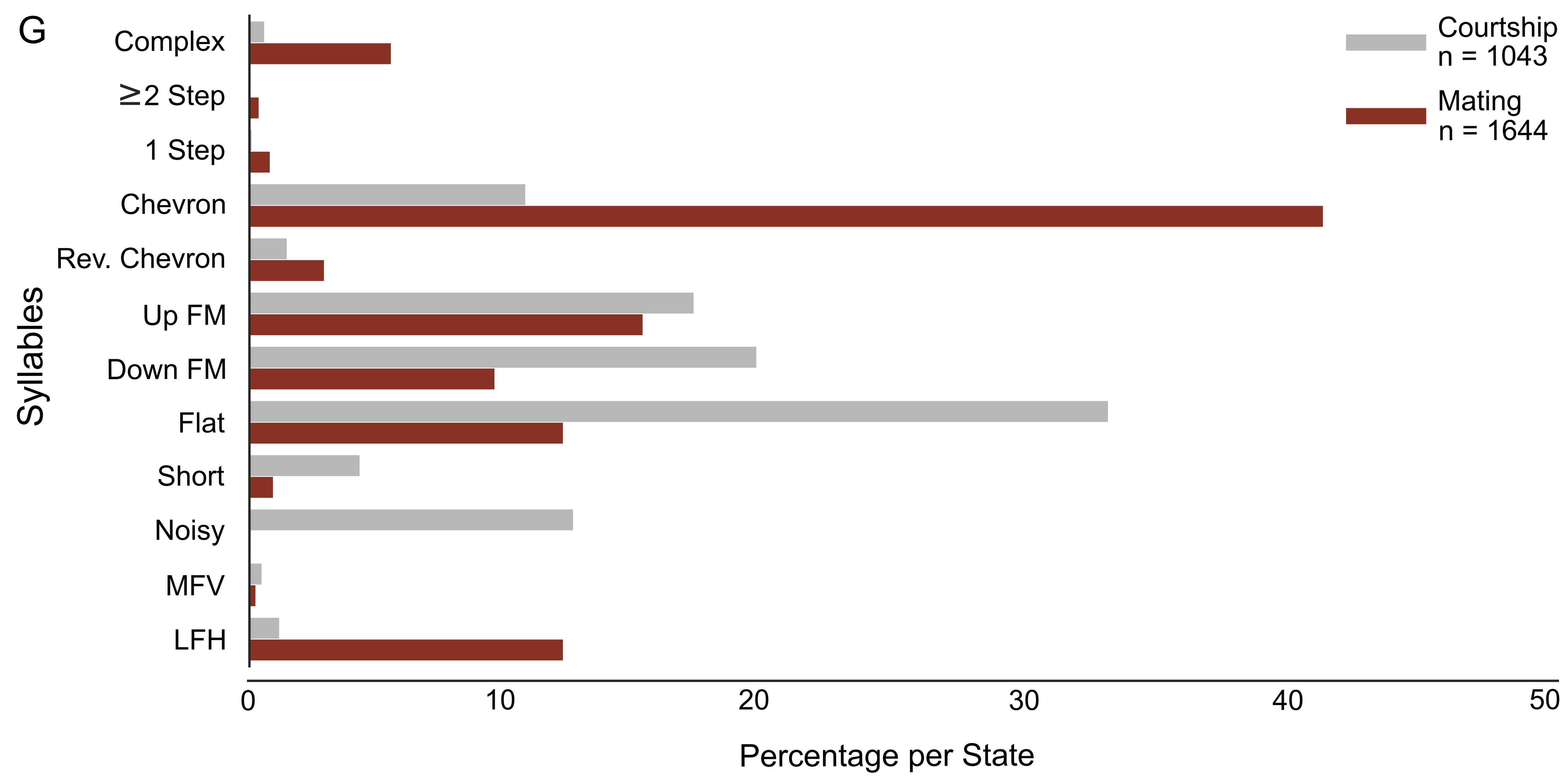

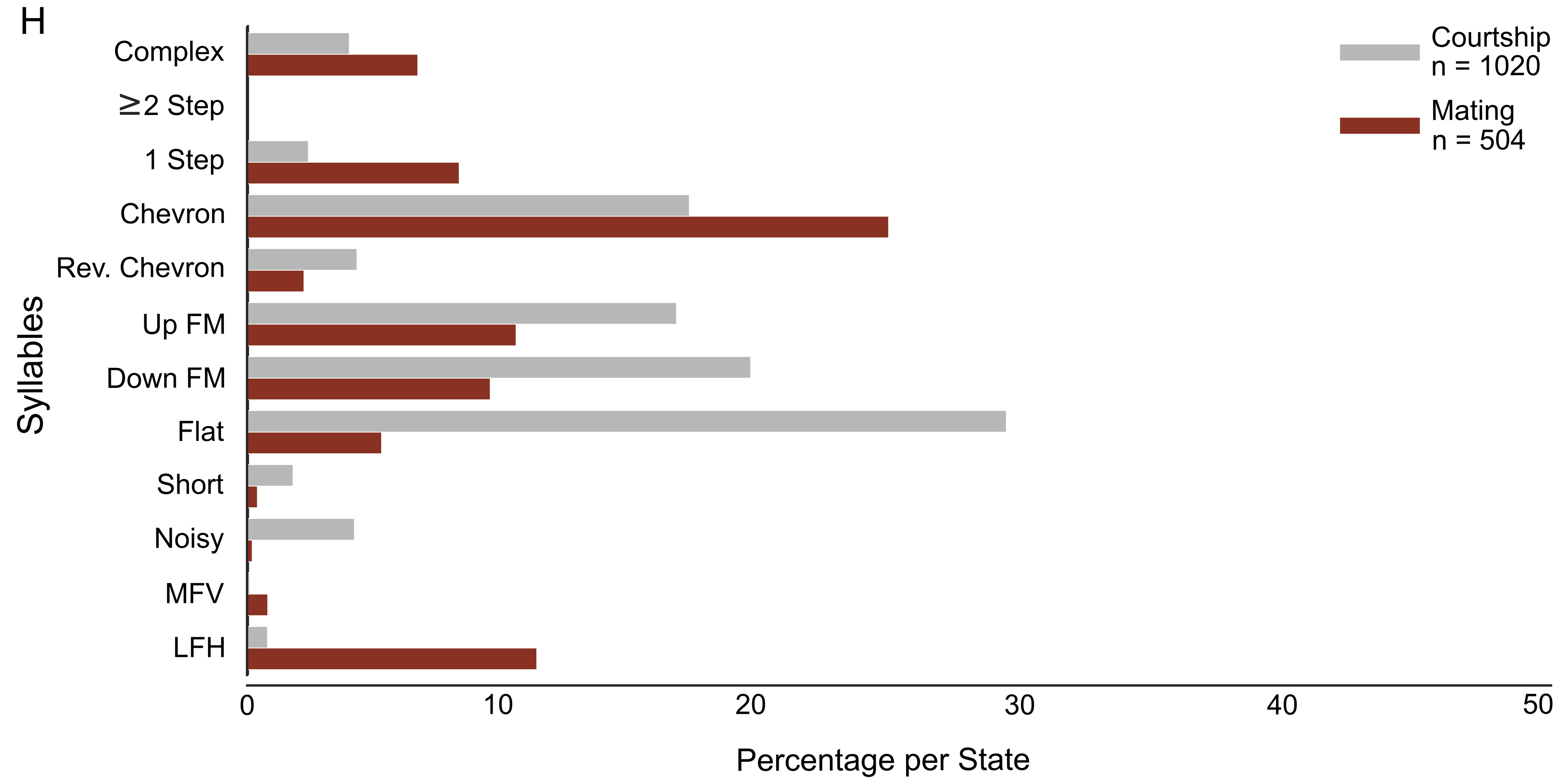
