## Supplementary material for "Acoustic features of and behavioral responses to emotionally intense mouse vocalizations": Figure caption- Supplementary Figure 1

**Supporting Figure 1. Syllable composition change for each pair of male-female with change of interaction intensity. A-H.** Percentages of each syllable type recorded across courtship (gray) and mating (red) interactions for each pair of male-female included in vocalization recordings for Experiment I. The total number of syllables for each context is indicated at the top right corner of each panel (total syllables in courtship, n=6056; mating, n=7237 syllables).
